# St18 specifies globus pallidus projection neuron identity in MGE lineage

**DOI:** 10.1101/2021.08.03.454907

**Authors:** Luke F. Nunnelly, Melissa Campbell, Dylan I. Lee, Guoqiang Gu, Vilas Menon, Edmund Au

**Affiliations:** Department of Pathology & Cell Biology, Columbia University Irving Medical Center, New York NY, 10032, USA; Department of Neurology, Columbia University Irving Medical Center, New York NY, 10032, USA; Department of Cell and Developmental Biology, Vanderbilt University School of Medicine, Nashville TN, 37232, USA; Department of Rehabilitation and Regenerative Medicine, Columbia University Irving Medical Center, New York NY 10032, USA; Columbia Translational Neuroscience Initiative Scholar, Columbia University Irving Medical Center, New York NY, 10032, USA

## Abstract

The medial ganglionic eminence (MGE) is a progenitor domain in the subpallium that produces both locally-projecting interneurons which undergo tangential migration in structures such as the cortex as well as long-range projection neurons that occupy subcortical nuclei. Very little is known about the transcriptional mechanisms specifying the migratory behavior and axonal projection patterns of these two broad classes of MGE-derived neurons. In this study, we identify St18 as a novel transcriptional determinant specifying projection neuron fate in the MGE lineage. St18 is transiently expressed in the MGE subventricular zone (SVZ) and mantle, and we assessed its function using an ES cell-based model of MGE development. Induction of St18 is sufficient to direct ES-derived MGE neurons to adopt a projection neuron-like identity as defined by migration and morphology. Using genetic loss-of-function in mice, we find that St18 is required for the production of globus pallidus pars externa (GPe) prototypic projection neurons. Single cell RNA sequencing revealed that *St18* regulates MGE output of specific neuronal populations: in the absence of St18, we observe a large expansion of cortical interneurons at the expense of putative GPe neurons. Through gene expression analysis we identified a downstream effector of St18, Cbx7, which is a component of Polycomb repressor complex 1. We find that Cbx7 is essential for projection neuron-like migration and is not involved in St18-mediated projection neuron-like morphology. Our results characterize a novel transcriptional determinant that directs GPe prototypic projection neuron identity. Further, we identified a downstream target of St18, Cbx7, which regulates only the migratory behavior of long-range projection neurons, suggesting that specific features of MGE projection neuron identity may be governed in a compartmentalized fashion by distinct transcriptional modules downstream of St18.

## Introduction

The medial ganglionic eminence (MGE) is a neurogenic progenitor zone that gives rise to both short-range interneurons and long-range projection neurons that distribute broadly throughout the brain^1–3^. Interneuron versus projection neuron is one of the fundamental organizing principles of nervous system architecture, representing local versus point-to-point flows of information, respectively. However, the means by which multipotential neural stem cells in the MGE, and indeed elsewhere in the vertebrate nervous system, delineate interneurons from projection neuron cell fate is poorly understood. The MGE is best known for generating the majority of the GABAergic interneurons in the cortex, but it also produces inhibitory neurons that populate the hippocampus and striatum as well as subcortical nuclei, including the globus pallidus pars externa (GPe) and medial amygdala (MeA)^1–6^. MGE neural progenitors are defined by their expression of the transcription factor Nkx2-1^7–9^. However, as neural progenitors progress through and shortly after they exit the cell cycle, recent single cell RNA-seq (scRNAseq) studies have found that immature MGE neurons begin to express distinct gene repertoires en route to terminal differentiation^10, 11^. A key challenge is to link the expression of these early transcriptional antecedents with mature neuronal identity.

Two of the cardinal features of neuronal specification in the central nervous system are migratory behavior and the type of axon projections. During development, MGE neurons adopt distinct migratory routes. Cortical interneurons, for instance, migrate long distances through a process known as tangential migration in order to reach the neocortex^12^. MGE-derived projection neurons, by contrast, need only migrate short distances from their subpallial site of origin to contribute to subcortical nuclei including the basal ganglia^5^. Once neurons arrive in their appropriate location, interneurons and projection neurons elaborate morphologies appropriate for their function: Interneuron axonal and dendrite arbors project and branch locally, while the axons of projection neurons extend long distances.

The MGE generates subcortical projection neuronal lineages that contribute to the MeA and GPe. While much is known about MGE production of cortical interneurons^12^, the process by which MGE progenitors generate its various projection neuron lineages is still poorly understood. In the GPe, the majority of MGE-derived neurons are called prototypic neurons that express parvalbumin (PV) and primarily project caudally to the subthalamic nucleus (STN). They constitute a key link in the indirect pathway of the basal ganglia^13, 14^. In this circuit, the GPe receives inhibitory input from the striatum and projects to the STN where it delivers high frequency, tonic inhibitory tone^15–17^.

A set of factors that could potentially regulate MGE development is the Myelin Transcription Factors (Myt TFs) gene family. They include three paralogs, Myt1 (Nzf2), Myt1L (Myt2 or Nzf1), and St18 (Nzf3 or Myt3) that are highly expressed in the progenitor cells of the neural and neuroendocrine lineages^18–20^. They can either activate or repress transcription depending on cellular context^21^. In humans, Myt TF mutations have been associated with cancer^22^, neuronal dysfunction^23, 24^, and neuroendocrine abnormalities^25, 26^. In model organisms, Myt1 and Myt1L were reported to regulate endocrine/neuronal differentiation or trans-differentiation by repressing non-neuronal genes^19, 27^, while various MyT family members, including St18 regulate pancreatic islet cell function and survival^28–30^. The function of Myt1L and St18 *in vivo* during neural differentiation, however, remains largely unexplored.

Using an ES cell-based model of MGE development^31^, we found that St18 directs two aspects of projection neuron identity: migratory behavior and axonal morphology. Genetic loss of function of *St18* reveals a specific reduction in GPe prototypic neurons, a projection neuron population originating from the MGE. scRNA-seq analysis of the MGE indicated that *St18* regulates progenitor cell specification, which manifests as altered neuronal output. Finally, we identified Cbx7 as a downstream effector of *St18*, which regulates migration, while sparing *St18*-mediated projection neuron-like morphology suggesting that St18-mediated neuronal subtype specification is achieved through different downstream molecular modules independently regulating migration and axonal morphogenesis.

## Results

### *St18* is transiently expressed in the medial ganglionic eminence in the developing embryonic brain

St18 is a transcription factor transiently expressed in neural progenitors and immature neurons during embryonic development, including expression in the MGE^32^; Allen Institute, Developing Mouse Brain Atlas. We first confirmed embryonic expression of St18 by RNAscope. At E13.5, we found that St18 transcript is present in the MGE subventricular zone (SVZ) and mantle as well as in the marginal zone of the cortical plate (CP) (Figure 1A). We next investigated St18 expression by immunohistochemistry. At E13.5, we found that St18 protein is present in the MGE SVZ and mantle and is co-localized with MGE lineage neurons fate-mapped with a conditional tdTomato reporter (*Nkx2-1*^Cre^; Ai9) (Figure 1B-C) (Xu et al., 2008) but was largely absent in tangentially migrating MGE interneurons (Figure 1B, C). As with RNAscope, St18 protein is also present in the cortical plate (Figure 1 B, C). To validate our antibody, we tested embryonic brain tissue from *St18* (-/-) (whole animal *St18* null) and MGE-specific conditional knockout mice and compared to WT positive controls. Whole tissue *St18* (-/-) animals were generated by crossing an *St18* conditional allele (cKO) to a germline driver line (*E2a*^Cre^), and MGE-specific cKO animals were generated by crossing the *St18* cKO allele to an MGE specific driver line (*Nkx2-1*^Cre^). Histology for St18 revealed that St18 signal was absent throughout *St18* (-/-) embryos (Figure S1A, C). These findings are consistent with previous work characterizing the *St18* conditional allele and antibody^29, 33^. Taken together, our data is consistent with previously reported in situ hybridization data^32^Allen Institute, Developing Mouse Brain Atlas, and indicates that the genetic knockout strategy to ablate *St18* works as anticipated.

**Figure 1.**
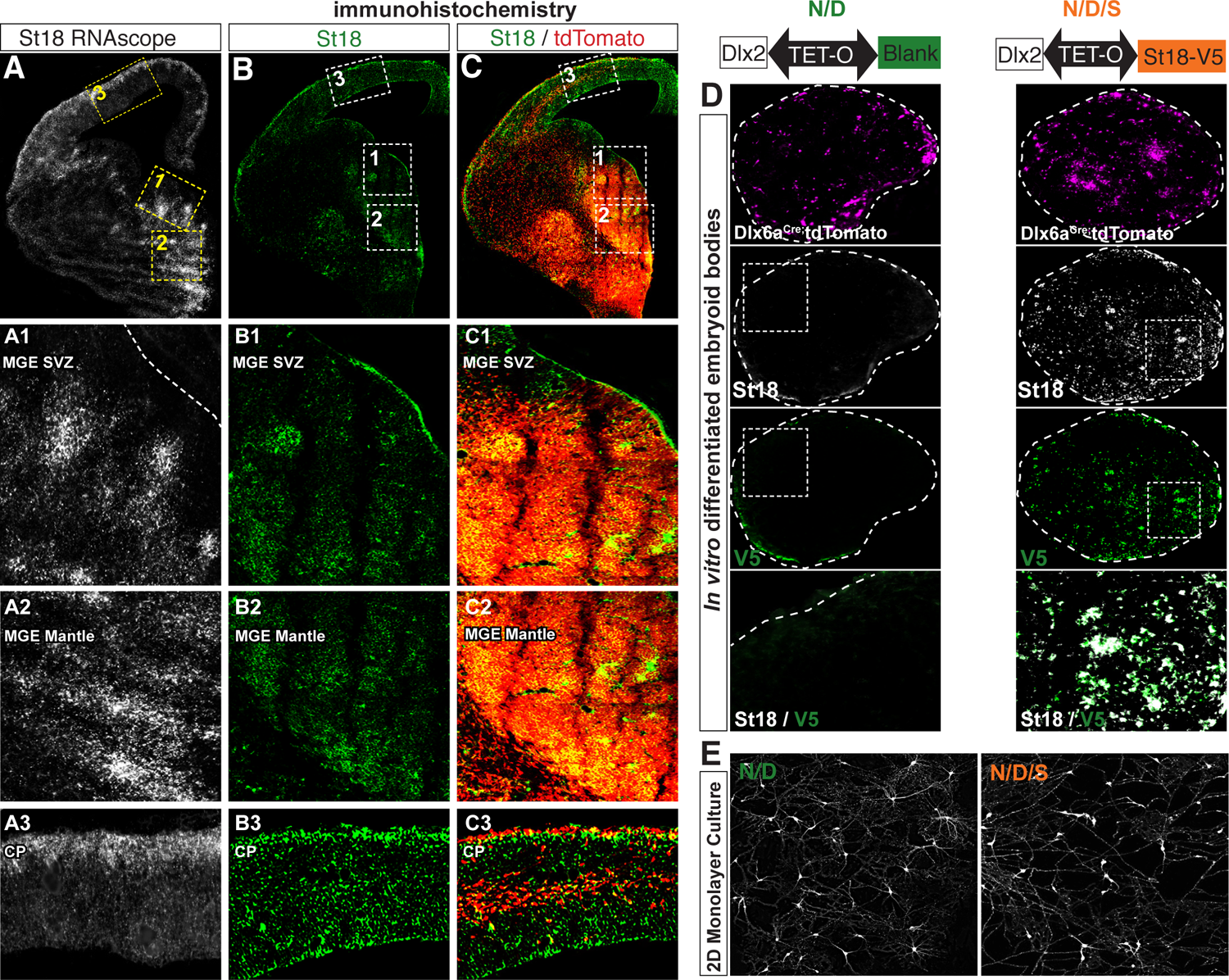
St18 expression in the in vivo MGE and St18 GOF in in vitro model of MGE development. (A) E13.5 WT embryo with RNAscope for St18 expression. Insets indicate higher power images of the MGE VZ (1) MGE Mantle (2) and CP (3). (B) E13.5 WT embryo with immunolabelled St18. Insets indicate higher power images of the MGE (1) GP (2) and CP (3). (C) E13.5 WT embryo with (B) merged with MGE lineage fate map (Nkx2-1Cre; Ai9). Subsets same as (B). (D) Schematic of transcriptional GOF in N/D and N/D/S conditions. N/D and N/D/S ES cell lines are differentiated over 12 days into EBs. Sectioned differentiated EBs from N/D and N/D/S shown below schematic with conditional fate-map (Dlx6aCre; Ai9) and immunolabeling for St18 and V5 tag. (E) 2D monolayer cultures from dissociated EBs shown with conditional fate-map (Dlx6aCre; Ai9). EBs are gently dissociated and then plated at high density to optimal survival and then cultured for 12 days before imaging.

### Transcriptional induction of St18 in ES cell-derived MGE progenitors alters their migratory and morphological characteristics consistent with projection neuron-like identity

To test the function of St18 in the MGE, we employed an *in vitro* ES cell differentiation model of MGE development, which we previously developed as a tool to systematically test transcription factors expressed in the MGE^31, 34^. Briefly, we start with a parental ES reporter line (Dlx6a-Cre; Ai9^35^) that fate maps subpallial neuronal lineages. The line is sequentially transcriptionally specified by Nkx2-1 and Dlx2. Nkx2-1 imparts MGE identity ^7, 9^ while Dlx2 further specifies GABAergic neuronal lineages^36–38^. The system makes use of a bidirectional Tet-O cassette that allows us to test candidate transcription factors in gain of function alongside Nkx2-1 and Dlx2 (Figure 1D). Using this system, we generated two lines: one control line for generic GABAergic MGE lineages using Nkx2-1 and Dlx2, which we named N/D, and a second line with Nkx2-1, Dlx2, and St18 (with a V5 tag), named N/D/S, to test how St18 impacts MGE neuronal output.

To verify St18 induction in the N/D/S line, we tested St18 and V5 expression by immunohistochemistry in N/D and N/D/S lines differentiated into embryoid bodies (EBs) and found co-localized labeling for St18 and V5 in the N/D/S line (Figure 1D). In contrast, St18 was detected only sparsely with an absence of V5 expression in N/D EBs (Figure 1D). Further, tdTomato induction was not grossly affected by St18 induction (Figure 1D). Next, we differentiated N/D and N/D/S EBs and dissociated them into monolayer cultures to examine tdTomato+ neurons. Here, we noticed a dramatic change: tdTomato fate-mapped N/D neurons were multipolar with complex morphologies reminiscent of interneurons, and consistent with our previous studies^31, 34^ while tdTomato+ N/D/S neurons appeared to have less complex morphologies and bore longer processes, similar to projection neurons (Figure 1E).

The MGE generates both short range interneurons as well as long range projection neurons^2, 3^. Our data indicated that St18 may direct MGE progenitor towards projection neuron identity at the expense of interneuron identity. To test this hypothesis, we more carefully examined N/D/S neurons and N/D neurons by two key features that delineate interneuron and projection neurons: migration and morphology. A defining characteristic of MGE-derived interneurons is their long-distance tangential migration from their birthplace in the ventral telencephalon into the cortex^39^. In contrast, MGE-derived projection neurons migrate shorter distances, where they remain subcortical and populate nuclei in the basal ganglia and amygdala^3–6, 40^ (Figure 2A). Thus, if St18 instructs MGE progenitors to become projection neurons, we would expect that N/D/S neurons will migrate less than N/D neurons. To test this, we embedded N/D and N/D/S EBs in Matrigel and measured the capacity of *Dlx6a*^Cre^; Ai9 fate mapped neurons to migrate out of the EB (Figure 2B). Using this assay, we found that N/D neurons robustly migrate out of EBs long distances radially into the Matrigel (Figure 2C). Conversely, N/D/S neurons migrate shorter distances than N/D controls (Figure 2D, E), oftentimes failing to leave the EB altogether (Figure 2D). Further, numerous N/D/S neurons that failed to migrate out bore long processes that emanated from the EB whereas migratory N/D neurons had leading unipolar and branched processes reminiscent of migratory interneurons (Figure 2C-D, inset 2).

**Figure 2.**
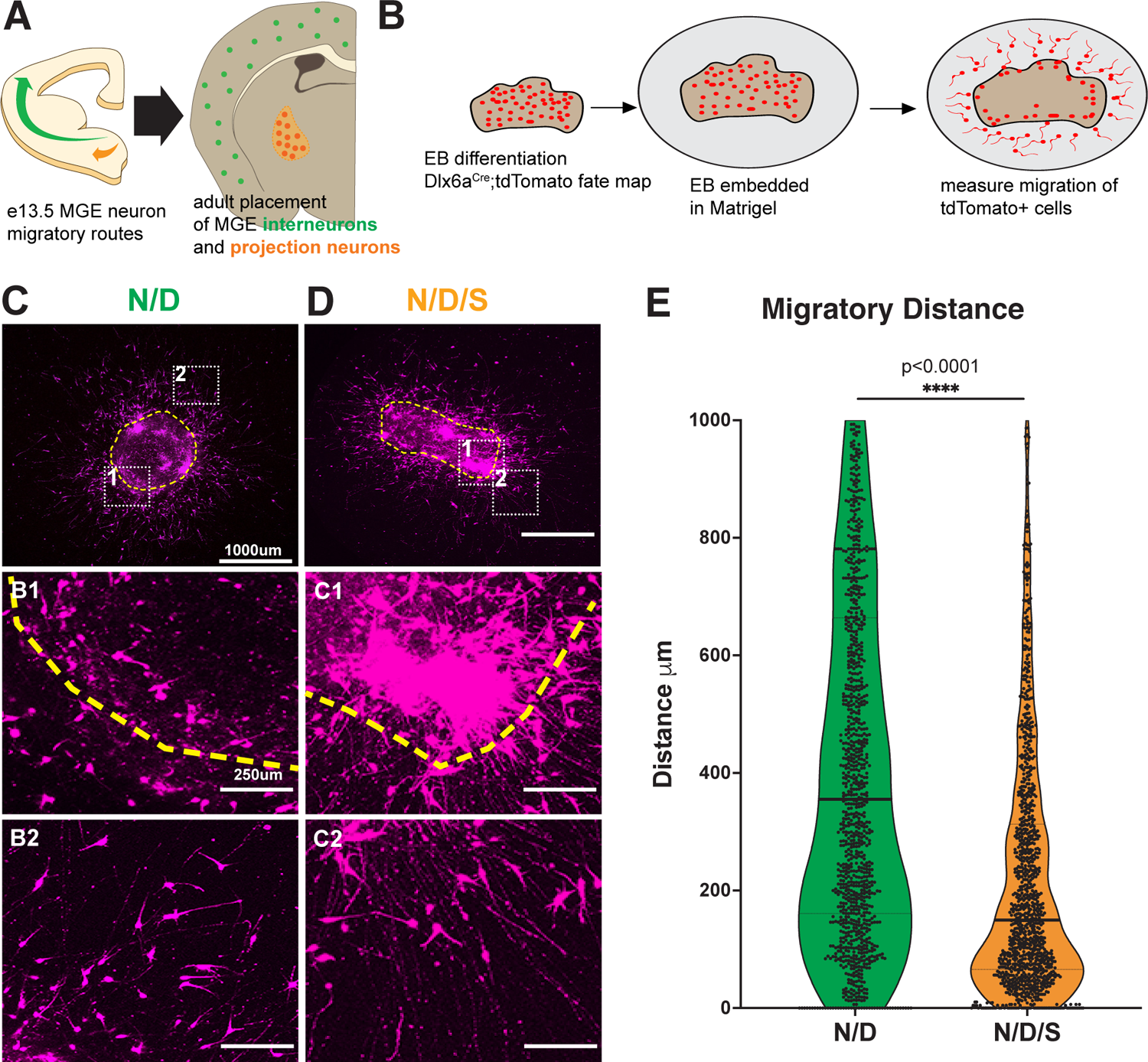
St18 GOF in vitro alters migratory behavior of MGE neurons. (A) Schematic representation (left) of MGE lineage neurons that adopt divergent migratory paths en route to (right) their final position in the adult brain. Cortical interneurons in green, globus pallidus neurons in orange. (B) Experimental design for Matrigel embedding experiments. N/D and N/D/S lines are differentiated into EBs for 12 days after which they are embedded in Matrigel. Fate-mapped neurons (Dlx6aCre; Ai9) are allowed to migrate radially into Matrigel for 4 days before quantification. (C) Representative image of N/D EB. Yellow dashed line indicates edge of EB. Insets show EB edge (1) and migrating fate-mapped neurons (2). (D) Representative image of N/D/S EB. Yellow dashed line indicates edge of EB. Subsets show neurons failing to migrate out of EB (1) and elaborated long processes (2). (E) Quantification of neuronal migration in (B) and (C): Migratory Distance (Mann-Whitney U Test). Dataset represents cells assayed from 6 EBs per condition; 1,137 N/D/S cells and 1,156 N/D cells.

This prompted us to examine the morphologies of N/D and N/D/S neurons in greater detail. To do so, we dissociated N/D and N/D/S EBs and plated them sparsely onto cortical feeders. This allows for long-term 2D culture of tdTomato+ neurons, which can be analyzed morphologically in isolation (Figure 3A). We found that N/D/S neurons bear far longer primary processes as well as longer average process length when compared with N/D neurons (Figure 3B, C, F). Next, we reconstructed N/D and N/D/S neurons and assayed morphological complexity by Sholl Analysis and found that N/D/S neurons are significantly less branched with longer, less complex processes compared to N/D neurons (Figure 3E, F; Additional morphological reconstructions in Figure S2). Thus, we find that N/D/S neurons are less migratory than N/D neurons and that N/D/S neurons exhibit a projection-like morphology (less-branched with longer processes) that are significantly different from the multipolar, shorter, complex morphologies of N/D neurons. Taken together, St18 also regulates neuronal migration and morphology and suggests that it is involved in specifying MGE projection neuron lineages.

**Figure 3.**
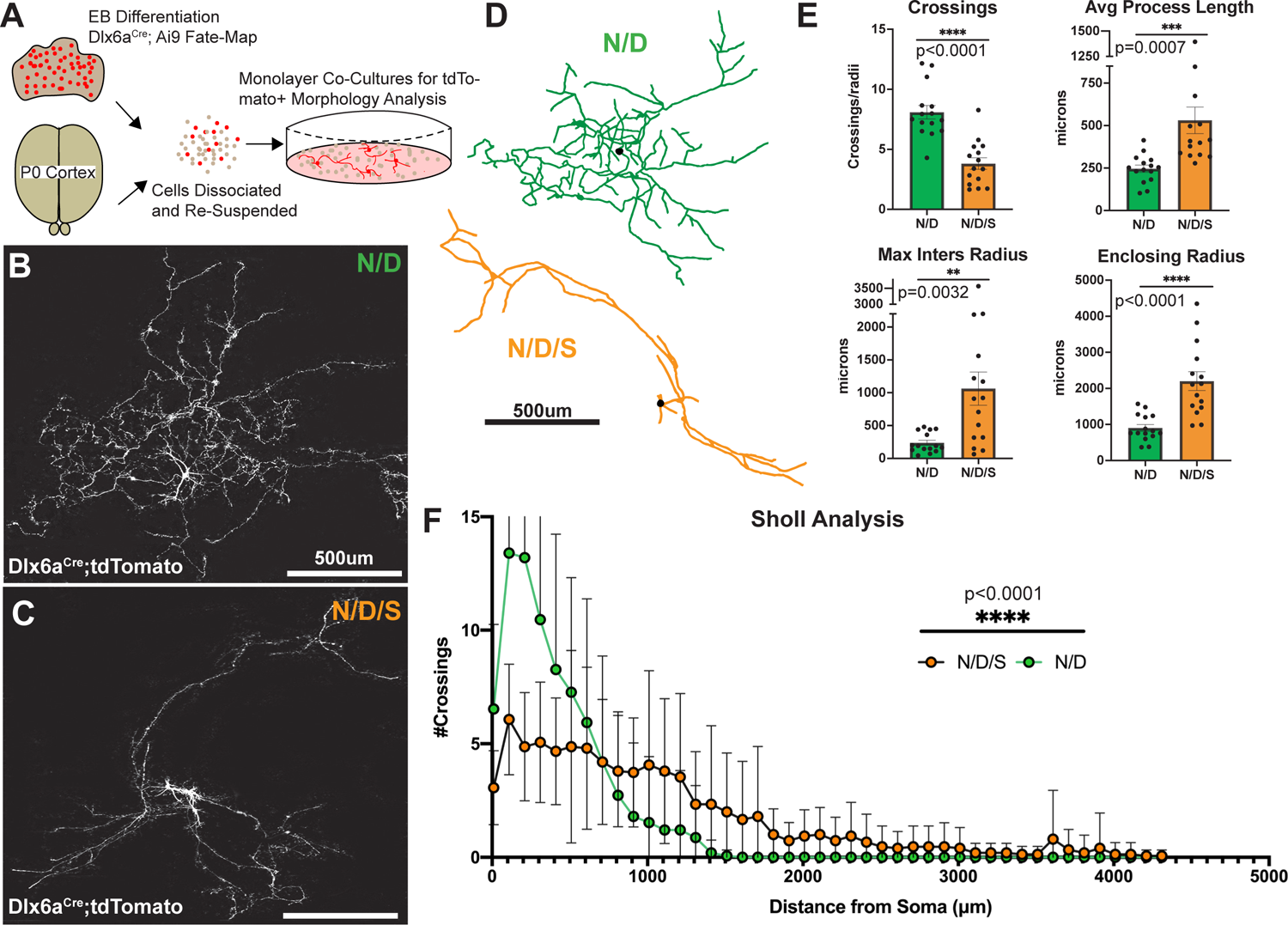
St18 GOF produces projection-like neuronal morphologies. (A) Experimental design of 2D monolayer tracing experiments. N/D and N/D/S lines are differentiated into EBs and dissociated into single cell suspensions. Concordantly, P0 corticies are dissociated and combined with low density EB suspensions for plating and whole cell tracing following fate-mapping (Dlx6aCre; Ai9). (B) Sample N/D neuron plated at low density with fate-mapping (Dlx6aCre; Ai9). (C) Sample N/D/S neuron plated at low density with fate-mapping (Dlx6aCre; Ai9). (D) Representative tracings of N/D and N/D/S neurons. (E) Sholl analysis curves for tracings in (D) and (E) (Two-Way ANOVA). (F) Sholl Crossings, Process Length, Maximum Intersection Radius, and Enclosing Radius quantification (Unpaired t-test). Dataset represents 15 reconstructed neurons per line.

### *St18* genetic loss of function results in specific loss of PV+ GPe prototypic projection neurons

Given our *in vitro* findings, we next investigated whether *St18* is necessary for the development of MGE projection neurons *in vivo.* We first tested if loss of St18 resulted in alterations in MGE proliferation and cell death by analyzing Ki67 and Caspase 3 in *St18* (-/-) embryos, respectively. No significant differences were observed in MGE cells for Ki67 or Caspase 3 in *St18* (-/-) embryos compared to WT controls (Figure S3a-b). We next tested whether *St18* (-/-) embryos exhibited defects in early MGE patterning Nkx2-1+ cells in either the MGE VZ/SVZ or ventrally in the developing globus pallidus (GP) (Flandin et al., 2010). We found that there is no difference in either the MGE or GP between *St18* (-/-) and WT embryos (Figure S3C). Together, we conclude that *St18* loss of function does not produce changes in proliferation, cell-death, or early MGE patterning. These findings are consistent with our observation that St18 expression is primarily found in SVZ and mantle regions.

The MGE produces GABAergic neurons that populate the cortex, hippocampus, striatum, medial amygdala (MeA), and globus pallidus pars externa (GPe)^3^. We surveyed the MGE neuronal lineage by immunohistochemistry, comparing *St18* (-/-) animals and littermate controls. We examined PV+ and SST+ neurons in the cortex, hippocampus, and striatum. We also tested the MGE component of the MeA, labeled by SST+ and FoxP2+^41–43^. In the GPe, we counted the number of MGE-derived prototypic neurons, determined by PV, Nkx2-1, and Er81 expression, the number of LGE-derived arkypallidal neurons, determined by Npas1 and FoxP2 expression, and GPe astrocytes, determined by S100b^5, 40, 44, 45^. We found that there was a significant decrease in PV+, Nkx2-1+, and Er81+ prototypic neurons in the GPe in *St18* (-/-) animals compared to controls (Figure 4). Importantly, we observed no significant difference in any other neural or glial component of the GPe, nor did we observe any significant difference in the other MGE derived neural lineages throughout the brain (Figure 4, Figure S4). Indeed, there were no changes in either the somatostatin (SST) or the PV neural populations in the cortex, hippocampus, or striatum following *St18* ablation, nor were there any changes in the SST and FoxP2 MGE derived neural lineages in the MeA (Figure S4). Thus, we only observe a specific reduction in PV+ GPe neurons in *St18* (-/-) animals.

**Figure 4.**
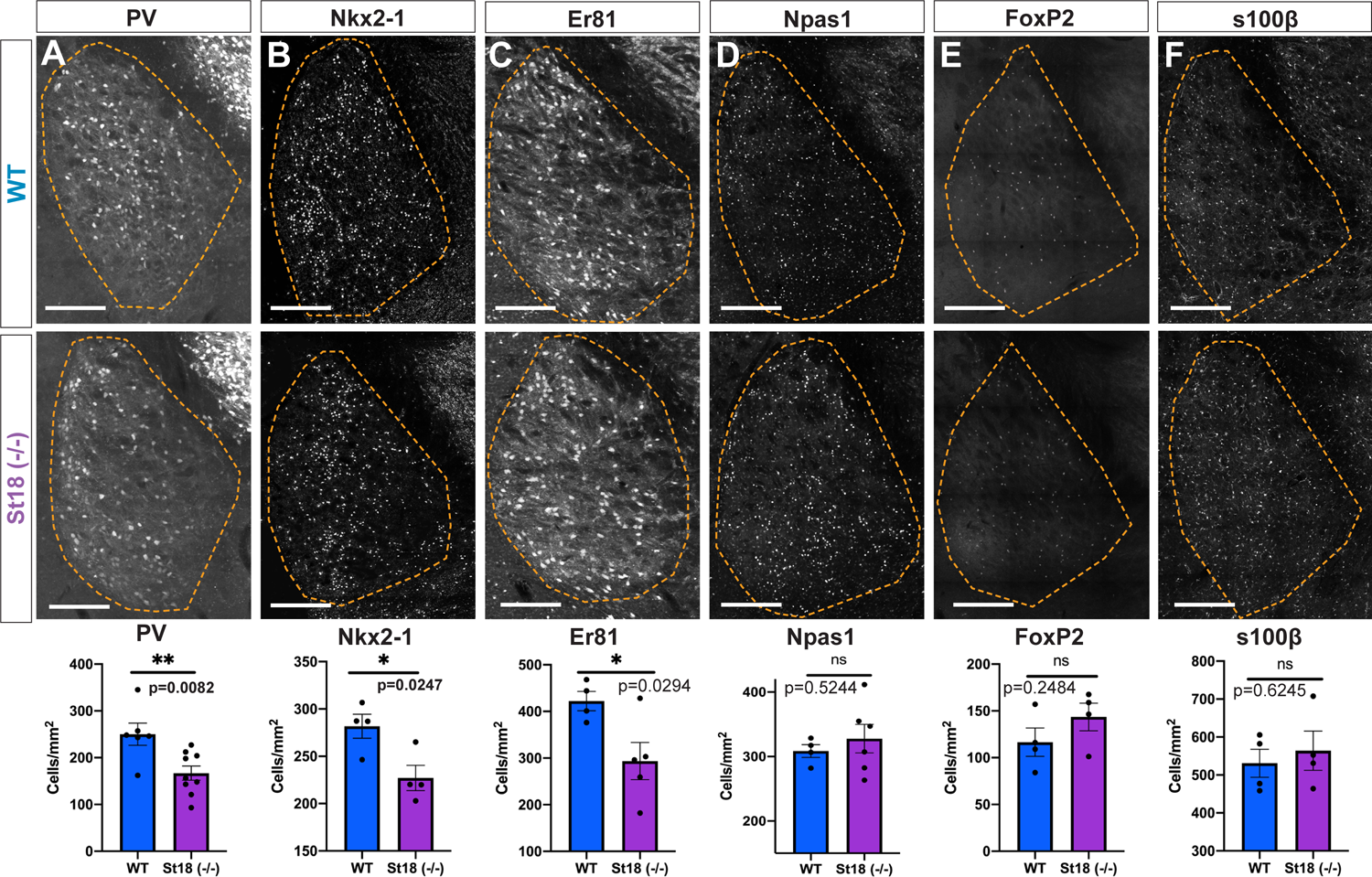
Whole animal St18 KO produces a specific loss of MGE lineage GPe projection neurons. Quantification of PV+ (A) Quantification of PV+ neurons (Unpaired t-test). N= 6 WT and 9 St18 (-/-). (B) Quantification of Nkx2-1+ neurons (Unpaired t-test). N= 4 WT and 4 St18 (-/-). (C) Quantification of Er81+ neurons (Unpaired t-test). N= 4 WT and 5 St18 (-/-). (D) Quantification of Npas1+ neurons (Unpaired t-test). N= 4 WT and 6 St18 (-/-). (E) Quantification of FoxP2+ neurons (Unpaired t-test). N= 4 WT and 4 St18 (-/-). (F) Quantification of s100B glia cells (Unpaired t-test). N= 4 WT and 4 St18 (-/-).

To investigate *St18* genetic ablation with greater resolution, we crossed the *St18* conditional allele to the MGE-specific driver line, *Nkx2-1*^Cre^ and the Ai9 conditional tdTomato reporter. This genetic strategy specifically ablates *St18* in the MGE while simultaneously fate-mapping the MGE neuronal lineage (*St18* cKO). We compared *St18* cKO (St18^fl/fl^; *Nkx2-1*^Cre^; Ai9) to *St18* heterozygote controls (St18^fl/+^; Nkx2-1^Cre^; Ai9). Efficient recombination of the St18 conditional allele in the MGE was confirmed by St18 immunohistochemistry (Figure S1B). We then quantified the number of fate-mapped tdTomato+ neurons throughout the brain to examine MGE lineages in *St18* cKO animals compared with controls. The number of tdTomato+ neurons in *St18* cKO animals was not significantly different in any region other than the GPe (Figure S5), where there was a significant decrease in tdTomato+ neuron density that was not due to a change in total GPe area (Figure S5A). Thus *St18* cKO, consistent with our findings in the *St18* (-/-), exhibits a specific loss of MGE lineage tdTomato+ fate-mapped neurons in the GPe.

We next repeated our histological analysis of the neural and glial components of the GPe. We found that, as in the *St18* (-/-) animal analysis, MGE derived prototypic neurons, identified by PV, Er81, and Nkx2-1 expression, were reduced while the other neuronal and glial components, determined by Npas1, FoxP2, and S100β, are spared in St18 cKO (Figure 5 A-C; Figure S6). Furthermore, like in *St18* (-/-) animals, *St18* cKO produces no significant change in the MGE derived neural lineages of the cortex, hippocampus, striatum, and MeA (Figure S7). Taken together, *St18* cKO phenocopies the *St18* (-/-), producing a specific loss of PV+ projection neurons in the GPe.

**Figure 5.**
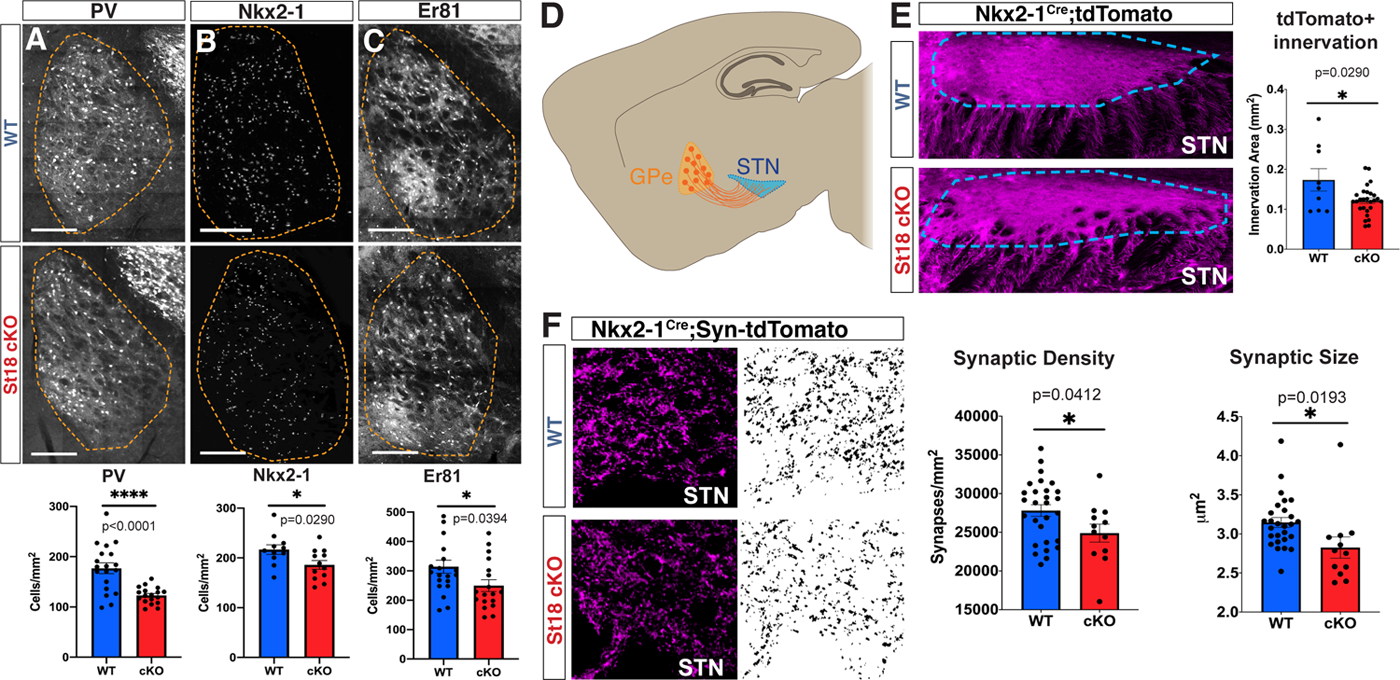
Conditional St18 KO produces a specific loss of MGE lineage GPe projection neurons. Quantification of PV+ (A) Quantification of PV+ neurons (Unpaired t-test). N= 19 WT and 18 St18 cKO. (B) Quantification of Nkx2-1+ neurons (Unpaired t-test). N= 11 WT and 12 St18 cKO. (C) Quantification of Er81+ neurons (Unpaired t-test). N= 20 WT and 18 St18 cKO. (D) Schematic representation of adult mouse brain in the parasagittal plane showing axonal projections of GPe prototypic neurons (orange) targeting the STN (blue). (E) WT and St18 cKO STN (outlined in blue) with MGE lineage whole cell labelled fate map (Nkx2-1Cre; Ai9). Quantification of axonal innervation area (Unpaired t-test). N= 9 WT and 27 St18 cKO. (F) WT and St18 cKO STN with (left) MGE lineage synaptophysin labelled fate map (Nkx2-1Cre; Ai34). tdTomato, (right) 3D segmented puncta quantified for puncta density and puncta size (Unpaired t-test). N= 27 WT and 12 St18 cKO.

### Disruptions in indirect pathway connections observed in St18 cKO mice

PV+ prototypic neurons in the GPe are an essential component of the basal ganglia indirect pathway. Prototypic neurons of the GPe project caudally to the subthalamic nucleus (STN) (Figure 5D), which then projects to motor thalamus and the spinal cord to regulate movement (Freeze et al., 2013). We tested whether loss of PV+ GPe neurons in St18 cKO animals resulted in a loss of innervation to the STN. The Ai9 reporter fills axonal processes, allowing us to examine projections from fate mapped prototypic GPe neurons to their target in the STN. We found a significant decrease in tdTomato+ axons in the STN in St18 cKO animals compared to controls (Figure 5E). To visualize synaptic innervation directly, we utilized a different reporter line, Ai34, that conditionally expresses synaptophysin-tdTomato (syn-tdT) upon Cre recombination. Syn-tdT puncta were segmented and quantified in the STN, revealing that puncta density and average puncta size were significantly reduced in *St18* cKO (Figure F). As a control, we examined Ai34 puncta density in the dorsal striatum (dStr) and the GPe, regions that are not primary targets of PV+ prototypic neurons in the GPe and found no significant differences (Figure S6D,E). Taken together, *St18* cKO results in reduced innervation of STN due to reduced axonal projections, synaptic density as well as reduced synaptic puncta size, all of which track with a reduction of PV+ prototypic neurons present in the GPe.

### scRNA-seq analysis of WT and St18 (-/-) MGE reveals specific alterations in neuronal output

Given the highly specific reduction in GPe prototypic projection neurons observed in *St18* mutants, we next examined how cell loss occurs. We found that overall proliferation, cell death and Nkx2-1+ progenitors were not significantly altered in the mutant MGE (Figure S3). We therefore explored the possibility that *St18* directs the cell fate of GPe prototypic neurons during progenitor cell specification. To do so, we performed single cell RNA-seq (scRNA-seq) on e14.5 *St18* (-/-) and WT littermate control MGE (n = 2 biological replicates per group; median of ∼9500 cells passing quality control per biological replicate). We chose to examine *St18* (-/-) instead of *St18* cKO MGE to guard against incomplete recombination with *Nkx2-1*^Cre^, which is unable to recombine in the MGE/LGE sulcus region^46^. We found that St18 transcript was still present in St18 (-/-) samples, although at significantly lower levels in neurons (Figure S10A; p=9.184×10^-5^; Wilcoxon rank sum test). This finding is consistent with previous work describing the *St18* conditional allele in that the transcript is still detectable after recombination, but at lower levels and likely reflects a degree of nonsense-mediated decay^29, 33^. Importantly, and consistent with previous work^29^, we found a lack of St18 protein signal in the *St18* (-/-) embryo by immunohistochemistry using a protein that recognizes the N-terminus (Figure S1). Thus, the N-terminal fragment of St18 that is encoded by the recombined St18 transcript is not detectable, likely due to protein instability.

Sequencing data was analyzed in a similar manner to a previous study of MGE progenitors and neurons^11^. After filtering cells with fewer than 200 genes detected, we performed a standard clustering analysis using the Seurat package in R^47^. Briefly, we identified high variance genes, followed by principal component analysis (retaining 49 PCs) and Louvain-based community detection clustering. Following subclustering of neurons and progenitors, we identified 13 progenitor cell clusters (3039 total cells in WT, 4018 total cells in *St18* (-/-)) and 21 neuronal clusters (5104 total cells in WT, 5904 total cells in *St18* (-/-)) (Figure 6A,B; Figure S9). A UMAP visualization of progenitor cell clusters in WT and *St18* (-/-) samples indicate that there are small differences in the relative proportions of clusters between the two groups, with a potential increase in progenitor clusters 7 and 9 in *St18* (-/-) (Figure 6A, C). However, a similar visualization of neuronal clusters shows more pronounced alterations in neuronal representation; indeed, statistical analysis using a Fisher exact test identified that the proportion of cluster 4 neurons, as a fraction of total neurons, was reduced 4.72-fold (p = 3.00 e^-103^), while the proportion of cluster 2 neurons was increased 4.74-fold (p = 1.39 e^-99^), and the proportion of cluster 5 neurons was increased 2.35-fold (p = 3.12 e^-28^) in *St18* (-/-) versus WT (Figure 6B, D). Analysis of gene expression in cluster 2 and 5 neurons revealed enrichment for cortical interneuron-related genes, for example: *Maf* (*c-Maf*)^47–50^, *Zeb2* (*Zfhx1b*, *Sip1*)^48, 51^, *EphA4*^52^, *Ackr3* (*CxcR7*)^53, 54^ and ErbB4^55^ over WT cluster 4 neurons (Figure S6E). St18 is not expressed at high levels in neuron clusters 2, 4 or 5 (Figure S10C). Thus, loss of *St18* likely affects progenitor cell production of neurons, resulting in a shift towards interneuron output (clusters 2 and 5) and away from cluster 4, which are likely to be nascent GPe prototypic neurons.

**Figure 6.**
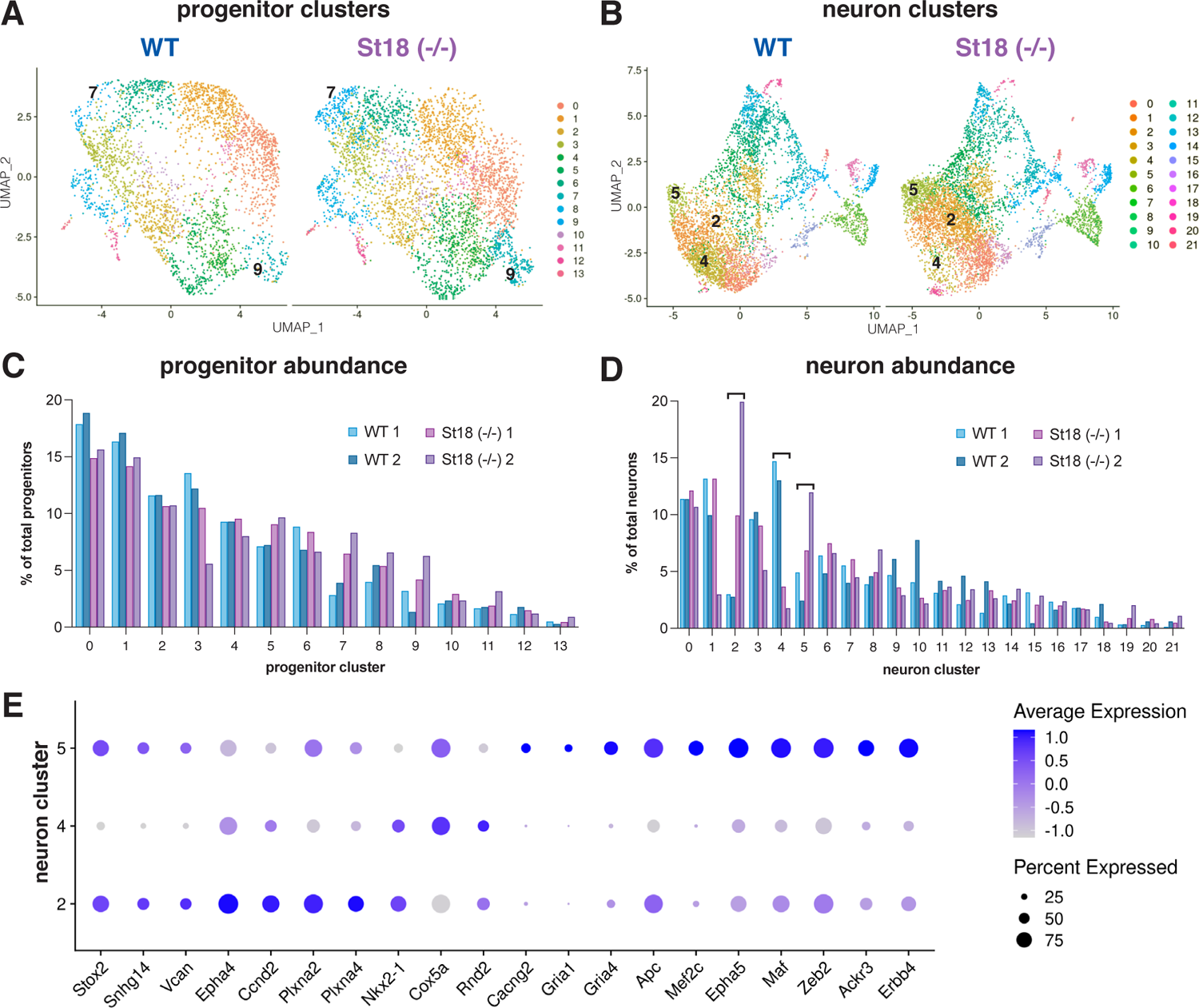
Single cell RNA-seq reveals that specific neuronal clusters are differentially produced upon St18 genetic ablation. (A) UMAP vizualization of 13 progenitor clusters identified in WT (left) and St18 (-/-) (St18 (-/-), right) e14.5 MGE. (B) UMAP visualization of 21 neuronal clusters identified in WT (left) and St18 (-/-) e14.5 MGE. Relative proportions of each (C) progenitor cluster and (D) neuronal clusters, biological replicates plotted separately. WT replicates 1 and 2 in shades of blue, St18 (-/-) replicates 1 and 2 in shades of purple. (E) Dot plot showing scaled average expression of differentially-expressed genes in neuron clusters 2, 4 and 5. Numbers indicated on UMAP visualizations in (A) and (B) denote clusters with the most pronounced alterations in proportional representation between WT and St18 (-/-). scRNA-seq dataset represents 2 biological replicates per genotype, 8,143 WT cells and 9,922 St18 (-/-) cells that passed quality control total.

### Cbx7 regulates projection neuron migration downstream of St18

To determine how St18 regulates projection neuron identity in MGE progenitors, we examined differential gene expression between WT and *St18* (-/-) MGE *in vivo* and then cross-referenced our findings with bulk RNA sequencing (RNAseq) in N/D and N/D/S neurons *in vitro*. We found that Cbx7 was one of the most differentially expressed genes in N/D/S line vs. N/D; it was strongly upregulated in the N/D/S line (log2FC = 2.09, ∼4.25-fold) (Figure 7A). We also found Cbx7 in our scRNA-seq analysis, but it was detected at overall low levels. However, there is a trend towards higher expression in cluster 4 neurons over cluster 2 in WT MGE; that trend was reversed in *St18* (-/-) MGE; since the proportion of cluster 4 neurons were significantly decreased in the *St18* (-/-) versus WT, it suggests a corresponding decrease in the representation of Cbx7-high neuronal signatures with *St18* genetic ablation.

**Figure 7.**
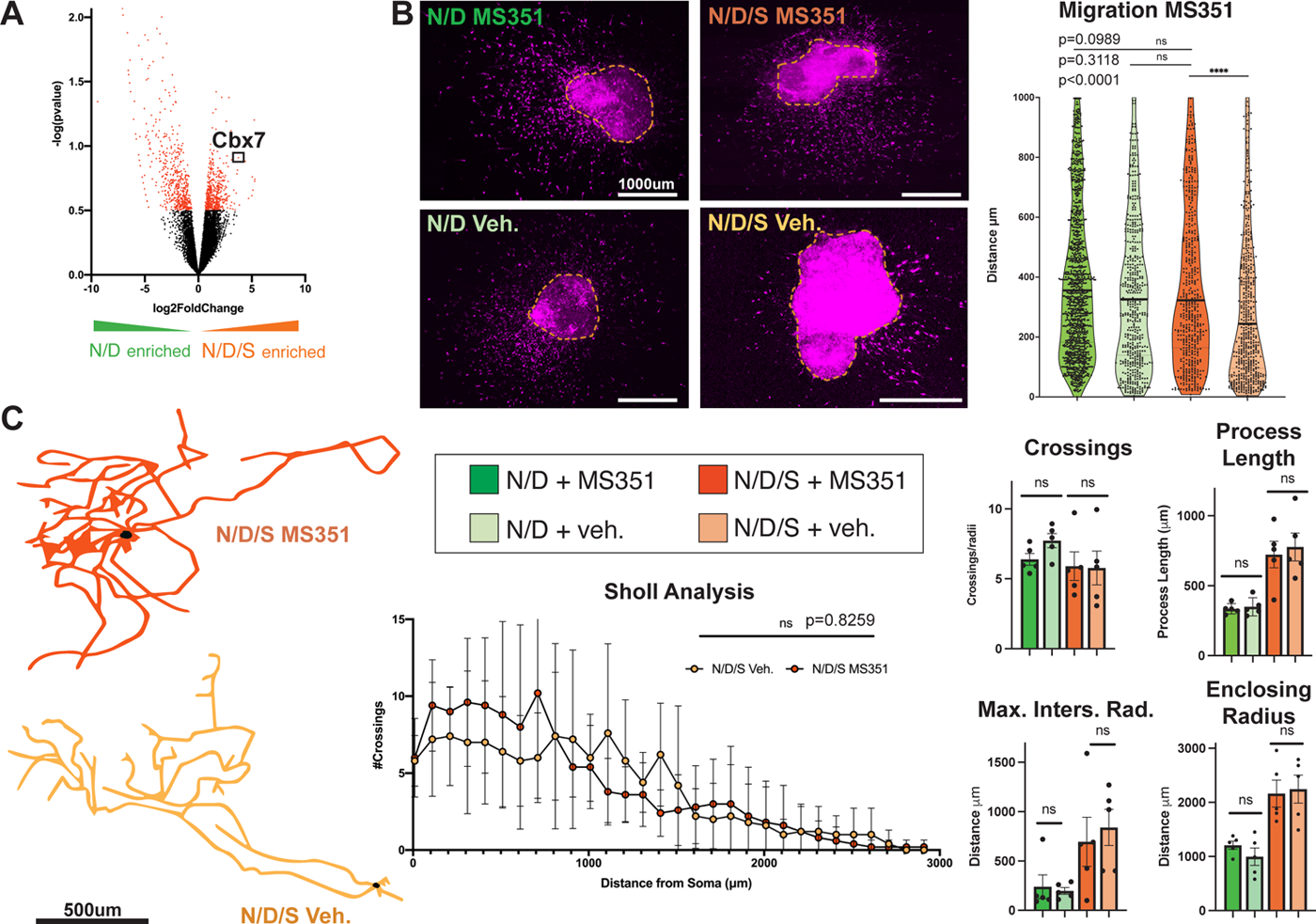
St18 regulates projection neuron migration but not morphology via upregulation of Cbx7. (A) MA plot of differentially expressed genes in N/D/S neurons compred to N/D neurons. (B) Sample images of Matrigel embedded N/D/S EBs treated either with MS351 or Veh. Inset shows higher power images. Arrowheads delinate migrating neurons in N/D/S MS351 panel and processes in N/D/S Veh. panel. Quantification to the right of the images (Mann-Whitney U Test). Dataset represents 3 EBs per condition; 531 N/D/S MS351 cells and 614 N/D/S Veh. cells and 1,301 N/D MS351 cells and 567 N/D Veh. cells. (C) Sample neuronal reconstructions in N/D/S MS351 and N/D/S Veh. conditions. Sholl analysis (2-way ANOVA) and Sholl statistics quantification to the right of the neuronal reconstructions (Unpaired t-test). N= 5 N/D/S MS351 and 5 N/D/S Veh. and 5 N/D MS351 and 5 N/D Veh.

Cbx7 is a component of the Polycomb repressor complex 1 (PRC1), which broadly enacts transcriptional repression^56^. Cbx7, via its association with the PRC1, is involved in both the early embryonic development of neural tissue^57^ as well as in post-mitotic differentiation of maturing neurons^58^, making it an attractive candidate to investigate as a potential downstream effector. We utilized a pharmacological inhibitor of Cbx7-mediated repression, MS351^59^ to test if it affected the migratory and morphological phenotypes of N/D/S neurons *in vitro*. To test the efficacy of MS351, we assayed P16^INK4a^ transcript levels by qPCR^60^ in N/D/S neurons since P16^INK4a^ transcription is negatively regulated by Cbx7-mediated PRC1 activity^61^. We found that N/D/S cells treated with MS351 had higher levels of P16^INK4a^ compared to N/D/S cells treated only with vehicle (ΔΔCt= −3.62; ∼150-fold increase MS351-treated vs. vehicle). We found that N/D neurons migrate significant further than N/D/S in our *in vitro* migration assay (Figure 2). However, following application of MS351, N/D/S neurons now migrate significantly further than vehicle-treated N/D/S neurons (Figure 7B). In fact, MS351-treated N/D/S neurons migrate as far as N/D neurons treated with MS351. Notably, MS351-treated N/D cell migration is not significantly different from vehicle-treated N/D neurons (Figure 7B). This result suggests that N/D/S neurons treated with MS351 lose projection neuron-like migratory behavior and instead, migrate in an interneuron-like manner. We next tested the effect of MS351 on neuronal morphology. Here, we found that MS351 had no effect on N/D/S or N/D morphologies as assessed by Sholl analysis (Figure 7C, Figure S11). Thus, Cbx7 regulates St18-mediated migration, while sparing St18-mediated projection neuron morphology altogether.

## Discussion

Many neural progenitors throughout the brain generate both interneurons and projection neurons, but how this crucial fate choice is determined is very poorly understood. In this work, we demonstrate that the transcription factor St18 directs MGE progenitors to adopt projection neuron identity at the expense of interneuron production. Using an *in vitro* assay, we find that St18 governs the migratory and morphological characteristics of ES-derived MGE neurons towards projection neuron identity, overriding the default interneuron fate of N/D cells. In whole animal null and conditional loss-of-function *St18* mutants, we observe a highly specific loss of prototypic GPe projection neurons, which is accompanied by a reduction in target innervation of the STN. By single cell RNA-seq analysis of *St18* (-/-) and wildtype MGE, we find that while MGE progenitors are largely unaffected, neuronal output is altered in *St18* mutants; specifically, we observe an expansion of nascent interneurons and a decrease in putative GPe prototypic neurons. Gene expression analysis identified Cbx7 as a downstream effector of St18. Through pharmacological inhibition, we show that Cbx7 governs St18-mediated cell migration, while St18-mediated projection neuron morphology is unaffected. We therefore describe the first transcriptional determinant in vertebrate brain that directs neural progenitors towards projection neuron over interneuron fate. Downstream of St18, Cbx7 specifically regulates migratory aspects of the projection neuron phenotype, demonstrating that different attributes of MGE projection neuron identity are parsed into distinct transcriptional programs.

Previous studies have identified genes responsible for various characteristics of MGE lineage neurons. These include specification of MGE ventricular progenitors by sustained expression of *Nkx2-1*^62, 63^, the regulation of PV cortical interneuron migration by *Sp8/Sp9*^64^, the differentiation and maturation of SST cortical interneurons by *Satb1*^65^, and the allocation of MGE lineage striatal neurons via chemoattractive and chemorepulsive cues in response to Nrg1/ErbB4 and EphB/ephrinB signaling, respectively^66^. St18, however, is the first identified transcription factor involved in the specification of projection neurons in the MGE lineage. Indeed, to our knowledge, St18 is the first transcription factor identified in the vertebrate brain that directs progenitor cells to delineate between projection neuron over interneuron identity.

St18 (also known as Myt3 and NZF3) is a member of the Myt family of transcription factors, characterized by a conserved zinc finger DNA binding motif^32^. Other Myt transcription factor superfamily members include Myt1, which promotes neurogenesis via inhibition of Notch signaling^19^, and Myt1L, one of the ‘BAM’ factors required to produce induced neurons ^67^ that drives neuronal fate by suppressing non-neuronal differentiation^27^. Further, Myt1 mutant mice exhibit defects in vagal nerve projections^20^. St18 has also been implicated in neuronal differentiation, acting in conjunction with Neurog1^68^, and is thought to act broadly through gene repression due to its high affinity DNA binding domain^69^. St18 also has a documented role in cancer, acting as a tumor suppressor^22^ - hence, its name: suppressor of tumorogenicity 18 - or as an oncogene^70^ depending on tissue type. St18 also mediates apoptosis in pancreatic beta cells^28^ and cell migration in islet cells through integrin signaling^71^. The latter is consistent with our finding that St18 regulates migration of MGE neurons in vitro.

Migratory patterns of MGE-lineage neurons are divergent. Cortical interneurons migrate long distances tangentially and then switch to radial migration in order to arealize throughout the cortex. Subpallial MGE neurons, on the other hand, migrate comparatively short distances to occupy the striatum, globus pallidus, and amygdala. While there is much known about factors that govern interneuron tangential migration into the cortex^72^, relatively little is known about short distance migration in subpallial MGE neurons. In fact, many of the known factors for MGE interneuron migration are guidance cue-related^72, 73^: Robo/Slit^74^, Eph/ephrin^52, 75^, semaphorin^62, 76^, Cxcr/Cxcl^53, 54^, RGMa-neogenin^77^, and Netrin^78^. Here, we find that St18 induction robustly curtails cell migration in N/D/S neurons versus N/D neurons. Given the paucity of guidance cues present in our *in vitro* migration assay, we surmise that St18 restricts the cell intrinsic capacity for MGE neurons to migrate long distances, which is a mechanism that is distinct from expression of guidance cues. During development, this restricted migratory capacity likely acts in concert with guidance cues in the orchestrated assembly of the GPe^5^ to ensure the appropriate placement of GPe prototypic neurons during development.

The GPe is comprised of two principle neuronal populations. One population are the arkypallidal cells that are primarily generated in the lateral ganglionic eminence and project back to the striatum. The second population are prototypic cells that are primarily MGE-derived and project mainly to the STN^2,5,40, 44^. Recent work describes seven distinct classes of GPe neurons^79^. Among these classes, four are PV+ prototypic cohorts, each with different, non-overlapping molecular marker expression and electrophysiological characteristics. Our analysis of St18 (-/-) and St18 cKO reveal a specific and significant loss of PV+ prototypic neurons accompanied by loss of Nkx2-1, Er81, and tdTomato + cells. We also observe a proportional reduction in fate-mapped synaptic innervation of the STN. Notably, we do not observe a complete loss of GPe prototypic neurons in St18 mutants, leading us to hypothesize that St18 is required for the specification of a subset of prototypic neurons and indicates that further work is required to parse the developmental origins of GPe cell types.

Two recent studies examined the dynamic process of MGE neurogenesis by scRNA-seq^10, 11^. They find that MGE progenitors share similar gene expression patterns and then rapidly diversify as they transition out of the cell cycle. Indeed, *St18* was identified as a marker gene for an MGE progenitor pool during this process of diversification^11^. However, shortly after MGE neurons are specified, they migrate extensively throughout the brain, making it a difficult task to link developmental gene expression with mature neuronal identity. In our scRNAseq analysis, we find a number of St18-expressing progenitor populations (progenitor clusters 0, 4, 7, 11 and 12; Figure S10B). However, we find that loss of *St18* does not largely affect the relative proportion of MGE progenitors. Instead, we observe sizable alterations in neuronal output, namely the loss of neuron cluster 4 and an expansion of neuron clusters 2 and 5. It is notable, however, that none of the affected neuronal clusters express St18 at high levels. This leads us to hypothesize that *St18* likely exerts its effect on cell identity in progenitors cells as they exit the cell cycle and begin to adopt neuronal identity. This has parallels with the transcription factor that defines MGE identity, Nkx2-1, which continues to influence cell fate up until cell cycle exit in a similar manner^7^.

We note that the specificity of the scRNA-seq phenotype mirrors the specific loss of GPe prototypic neurons in the *St18* mutant. Along this line, it is appealing to speculate that the identity of neuron cluster 4, which is strongly diminished in *St18* (-/-), are nascent GPe prototypic neurons. In support of this, we find that cluster 4 neurons are enriched in expression for *Nkx2-1*, whose sustained expression is found in subcortical MGE populations including GPe prototypic neurons^1,5^.

We also detect higher levels of *Rnd2* in cluster 4, which is associated with radial migration^80^. Given the close proximity of the MGE progenitor domain to the final location of the GPe, it is likely that immature GPe neurons migrate via radial versus tangential migration. In contrast, neuron clusters 2 and 5, which are expanded in *St18* (-/-), exhibit higher levels of *APC*, which is associated with tangential migration in cortical interneurons^81^. Indeed, we find numerous genes in clusters 2 and 5 that are reminiscent of cortical interneurons and express markers that indicate that they are nascent PV+ cortical interneurons: *Mef2C*^49^, and *Ccdn2*^82^. Thus, in the *St18* (-/-) MGE, we find selective alterations in three PV+ neuronal populations; given the reduction of cluster 4, accompanied by the expansions of cluster 2 and 5, we hypothesize that the 3 PV+ populations share a common lineal origin that is demarcated by the presence or absence of St18 expression at the progenitor cell stage.

While in *St18* (-/-) we observe robust expansions of neuron clusters 2 and 5, and similarly sharp reduction in neuron cluster 4 in our scRNA-seq analysis, it stands in contrast to the relatively moderate (∼35%) reduction in GPe prototypic neurons in *St18* mutants and the lack of an expanded cortical interneuron population. We hypothesize that selective survival and cell death helps normalize cell numbers postnatally, and accounts for the blunted impact observed in adult mutant animals. Certainly with cortical interneurons, a number of studies demonstrate that their cell numbers are under exquisite regulation^83–86^. It is conceivable that GPe neuronal numbers are similarly self-correcting, but since too few GPe prototypic neurons are produced in the St18 (-/-), selective survival alone is insufficient to bring cell numbers up to appropriate levels.

Very little is known about the molecular regulation of migration and axonal morphogenesis in neurons. To identify factors downstream of *St18* that regulate these two properties of projection neuron identity, we examined gene expression in *St18* (-/-) and wildtype MGE *in vivo* and cross-referenced with N/D versus N/D/S neuron gene expression *in vitro*. Through this process, we identified Cbx7. Cbx7 (chromobox 7) is a component of the Polycomb repressor complex 1 (PRC1). In this complex, it is involved both in early neural differentiation^57^ and post-mitotic neurite extension^58^. Cbx7 is also well-characterized in cancer. Mutations in *Cbx7* are correlated with increased glioblastoma multiforme tumor invasiveness^87, 88^. Restriction of migration is thought to be mediated by Cbx7 interaction with Wnt/b-catenin^89^ and the YAP-TAZ pathway^90^. Thus, the elevated levels of Cbx7 observed in decreased cancer metastasis are consistent with our findings that N/D/S neurons express higher levels of Cbx7 and migrate shorter distances.

We made use of a pharmacological inhibitor of Cbx7, MS351, that blocks Cbx7/PRC1-mediated H3K27Me3 transcriptional repression^59^ and found that it resulted in complete conversion of N/D/S such that they migrate like N/D neurons, while leaving St18-mediated morphology intact. This result raises the intriguing possibility that projection neuron cell fate is adopted in a piece-wise fashion, with migration and morphology being managed by separate transcriptional cascades. If so, this has parallels to what has been termed terminal selector genes in *C. elegans* and *Drosphila*. Terminal selectors are generally post-mitotic transcription factors responsible for directing or maintaining lineage characteristics^91^. For instance, mec-3 directs touch receptor identity in *C. elegans*^92^. In *Drosophila,* Apterous and Islet regulate morphological and physiological features^93, 94^. Indeed, some MGE genes such as *Tsc1* that regulates cortical interneuron intrinsic physiology^95^ also regulates specific aspects of interneuron identity. Our results suggest that *St18* and *Cbx7* may function in a similar arrangement, where the former sits atop a genetic hierarchy to direct cell fate while the latter governs gene repression that regulates the migratory portion of projection neuron identity. If this proves to be true, there may also be additional selector-like factors that govern projection neuron morphology and other aspects of GPe prototypic identity. Further work will be required to elucidate whether such an arrangement exists and whether this is a general organizing principle that regulates the cellular properties of other neuronal lineages arising from the MGE and throughout the brain.

---

**Figure S1.**
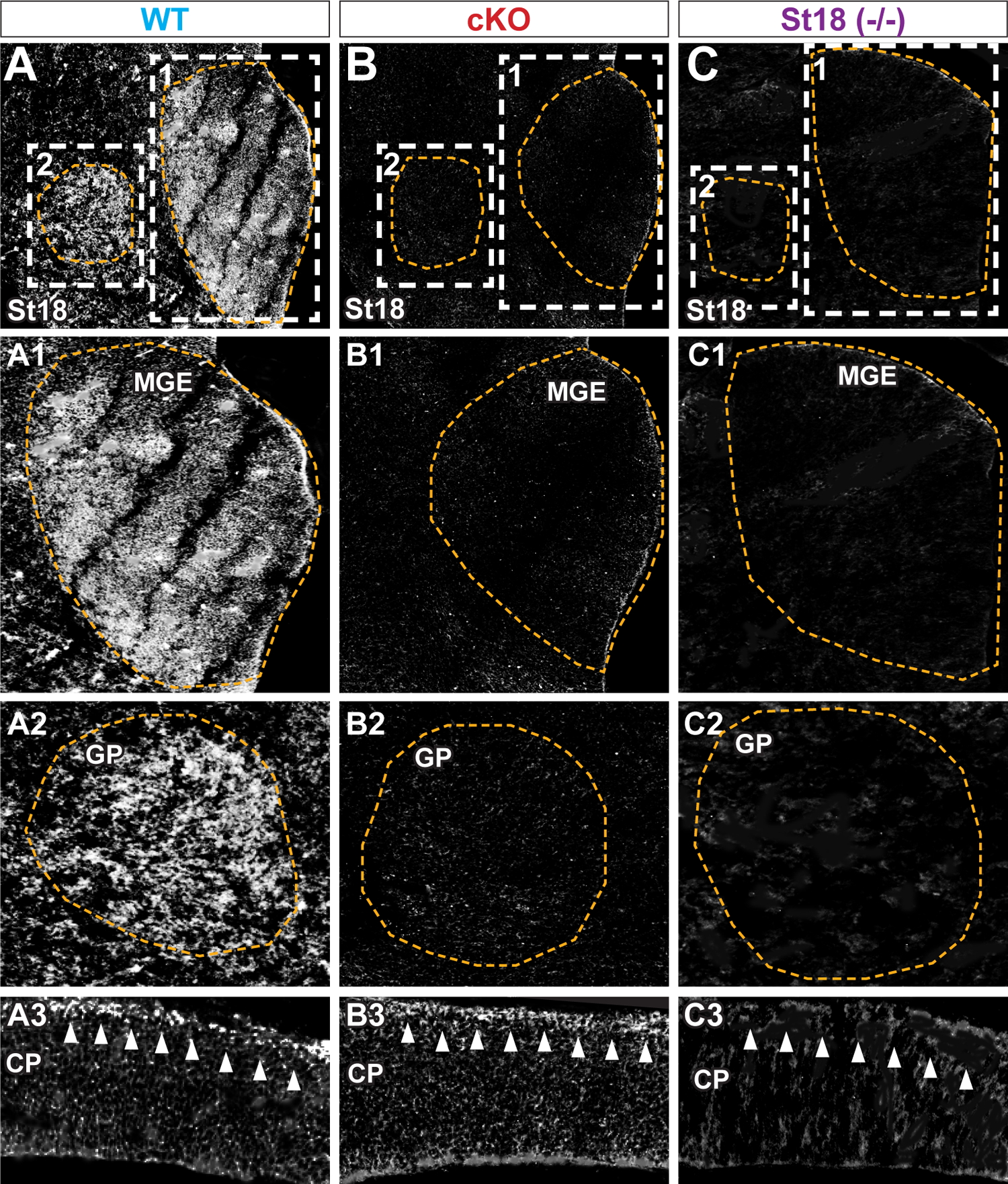
Validation of St18 antibody in WT, St18 cKO, and St18 (-/-) E13.5 embryonic MGE. (A) WT E13.5 MGE immunolabelled with St18 antibody. Insets indicate MGE (1) and GP (2). CP (3) shown below with arrowheads indicating location of St18 labelling. (B) St18 cKO E13.5 MGE immunolabelled with St18. Subsets are the same as (A). (C) St18 (-/-) E13.5 MGE immunolabelled with St18. Subsets are the same as (A).

**Figure S2.**
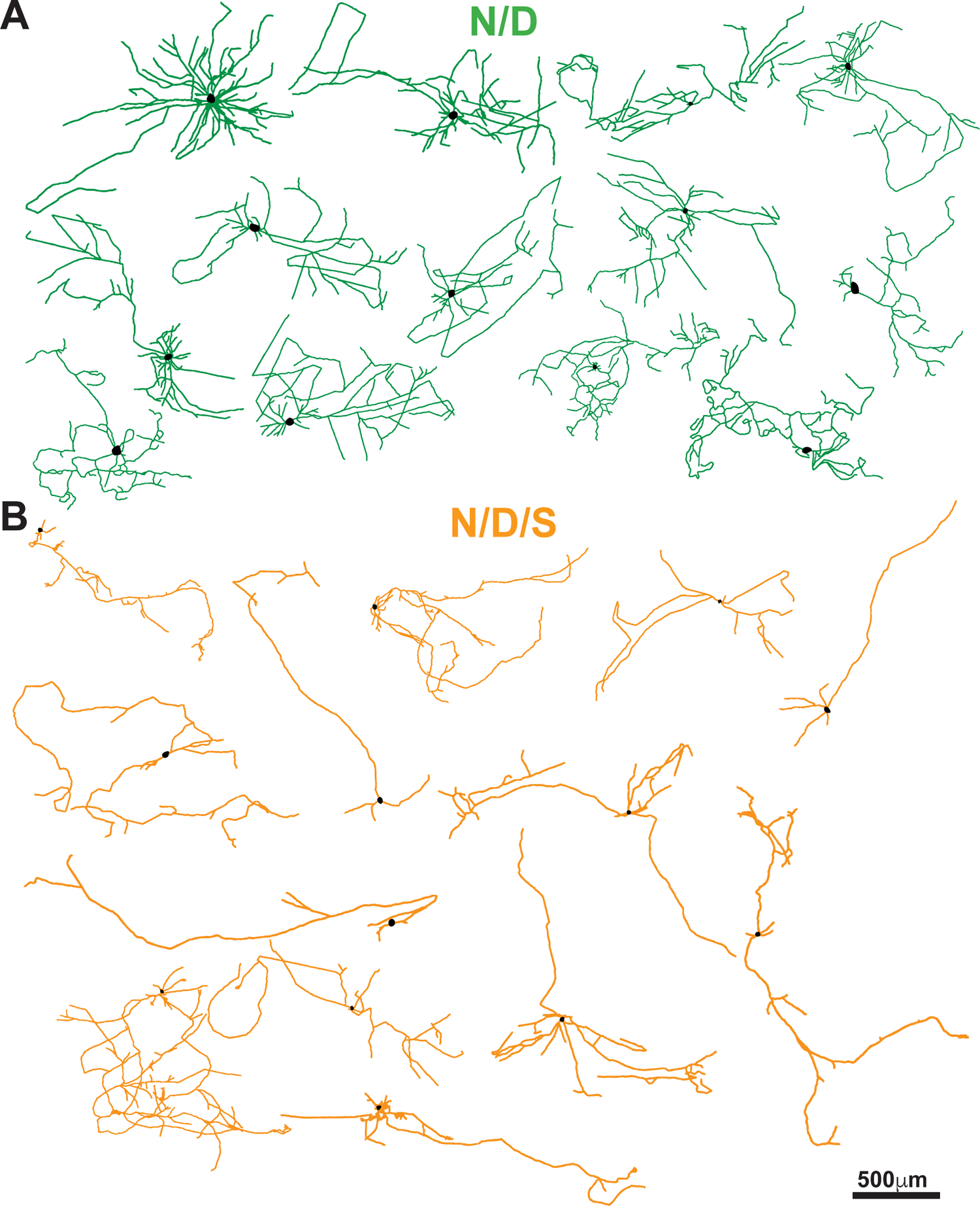
Additional morphological reconstructions of N/D and N/D/S neurons. Additional 2D tracings of ES-derived MGE neurons cultured in isolation on unlabeled cortical feeders. (A) N/D neurons in green. (B) N/D/S neurons in orange.

**Figure S3.**
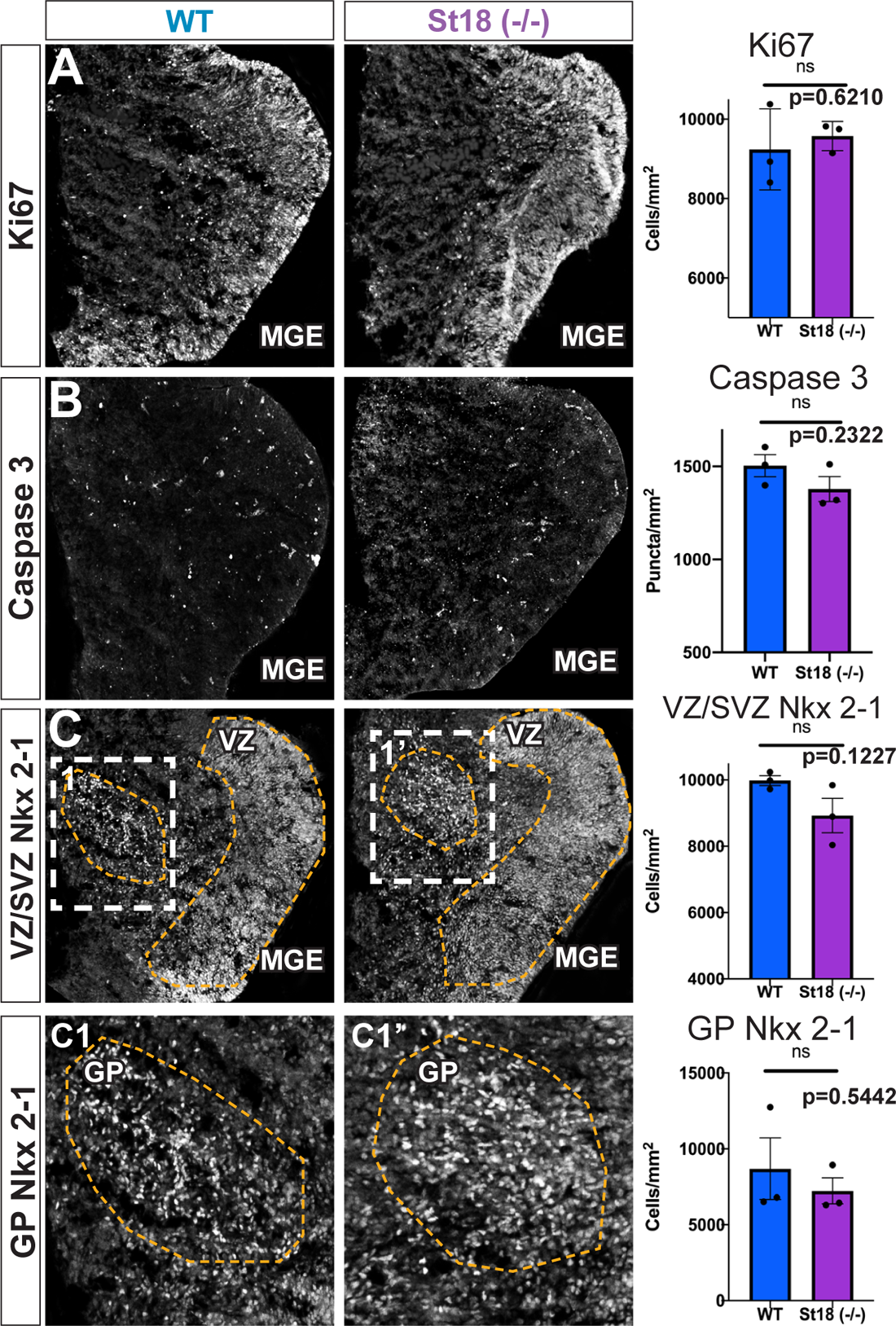
Whole animal St18 KO does not affect MGE proliferation, cell-death, nor MGE lineage specification at E13.5. (A) WT and St18 (-/-) E13.5 MGE with immunolabelled Ki67 (Unpaired t-test). (B) WT and St18 (-/-) E13.5 MGE with immunolabelled Casp3 (Unpaired t-test). (C) WT and St18 (-/-) E13.5 MGE with immunolabelled Nkx2-1. VZ is outlined in orange. Inset shows higher-powered imaged GP (Unpaired t-test). All datasets represent N= 3 per genotype.

**Figure S4.**
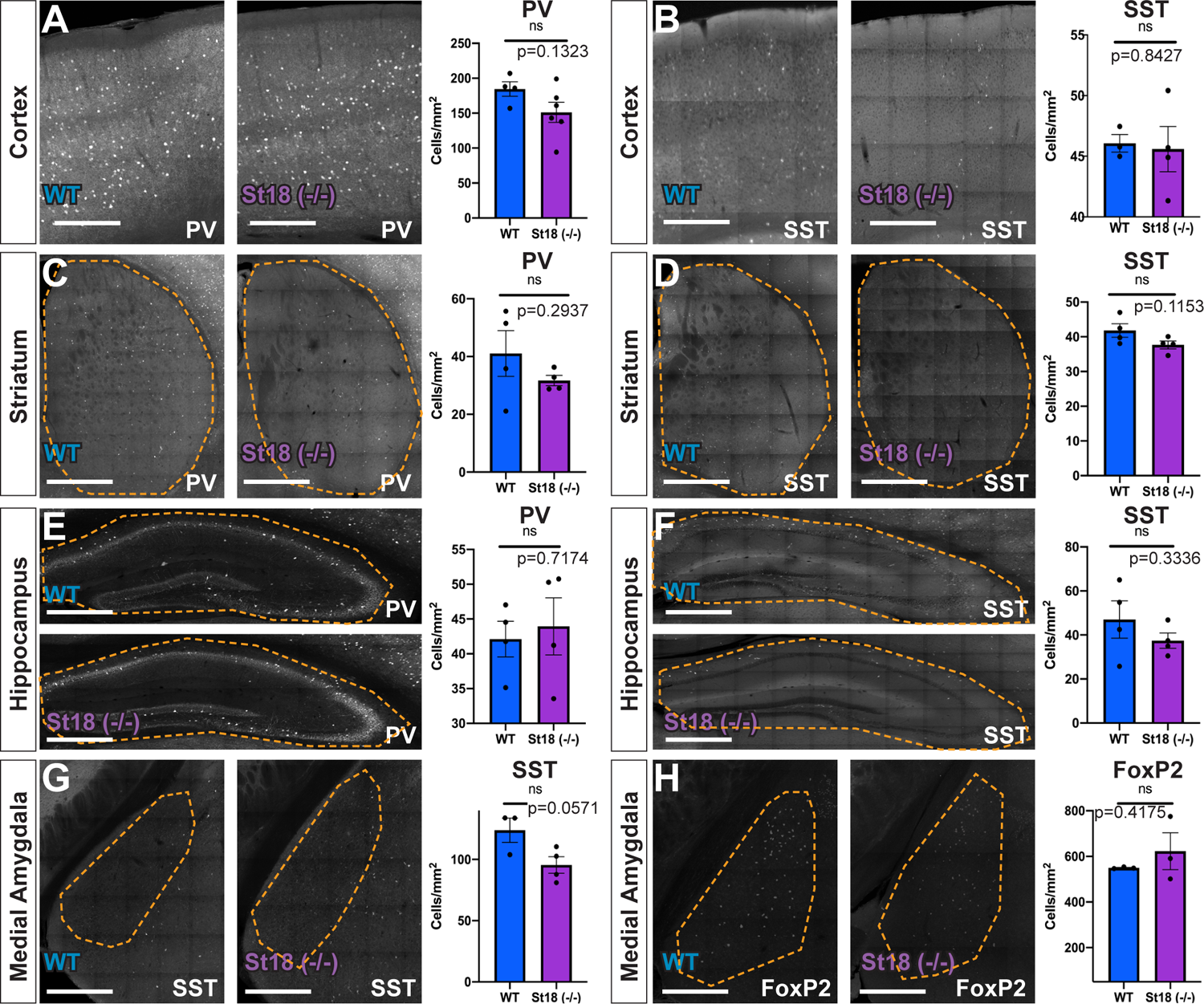
Whole animal St18 KO does not affect MGE lineages of the cortex, striatum, hippocampus, and MeA. Quantification of (A) PV+ (N= 4 WT and 6 St18 (-/-)) and (B) SST+ (N= 3 WT and 4 St18 (-/-)) neurons in the cortex. Quantification of (C) PV+ (N= 4 WT and 4 St18 (-/-)) and (D) SST+ (N= 4 WT and 4 St18 (-/-)) neurons in the striatum. Quantification of (E) PV+ (N= 4 WT and 4 St18 (-/-)) and (F) SST+ (N= 4 WT and 4 St18 (-/-)) neurons in the hippocampus. Quantification of (G) SST+ (N= 3 WT and 4 St18 (-/-)) and (H) FoxP2+ (N= 3 WT and St18 (-/-)) neurons in the MeA. (all data assessed by Unpaired t-test).

**Figure S5.**
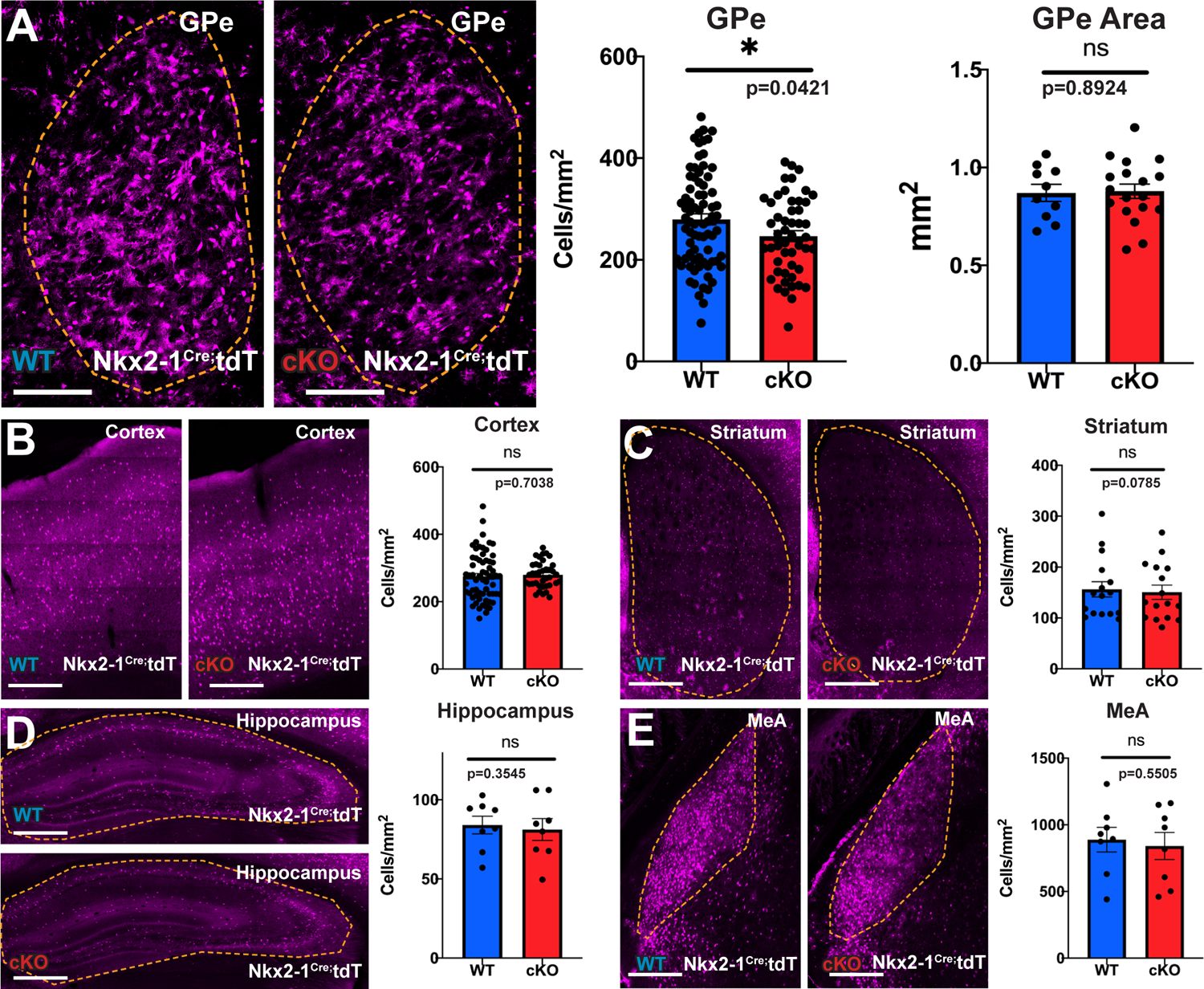
St18 conditional ablation in the MGE produces a specific loss of fate-mapped tdTomato+ neurons in the GPe. (A) WT and St18 cKO GPe with MGE lineage fate-map (Nkx2-1Cre; Ai9) (Unpaired t-test; N= 76 WT and 50 St18 cKO). (B) WT and St18 cKO Cortex with MGE lineage fate-map (Nkx2-1Cre; Ai9) (Unpaired t-test; N= 63 WT and 38 St18 cKO). (C) WT and St18 cKO Hippocampus with MGE lineage fate-map (Nkx2-1Cre; Ai9) (Unpaired t-test; N= 8 WT and 8 St18 cKO). (D) WT and St18 cKO MeA with MGE lineage fate-map (Nkx2-1Cre; Ai9) (Unpaired t-test; N= 8 WT and 8 St18 cKO). (E) WT and St18 cKO Striatum with MGE lineage fate-map (Nkx2-1Cre; Ai9) (Unpaired t-test; N= 16 WT and 16 St18 cKO).

**Figure S6.**
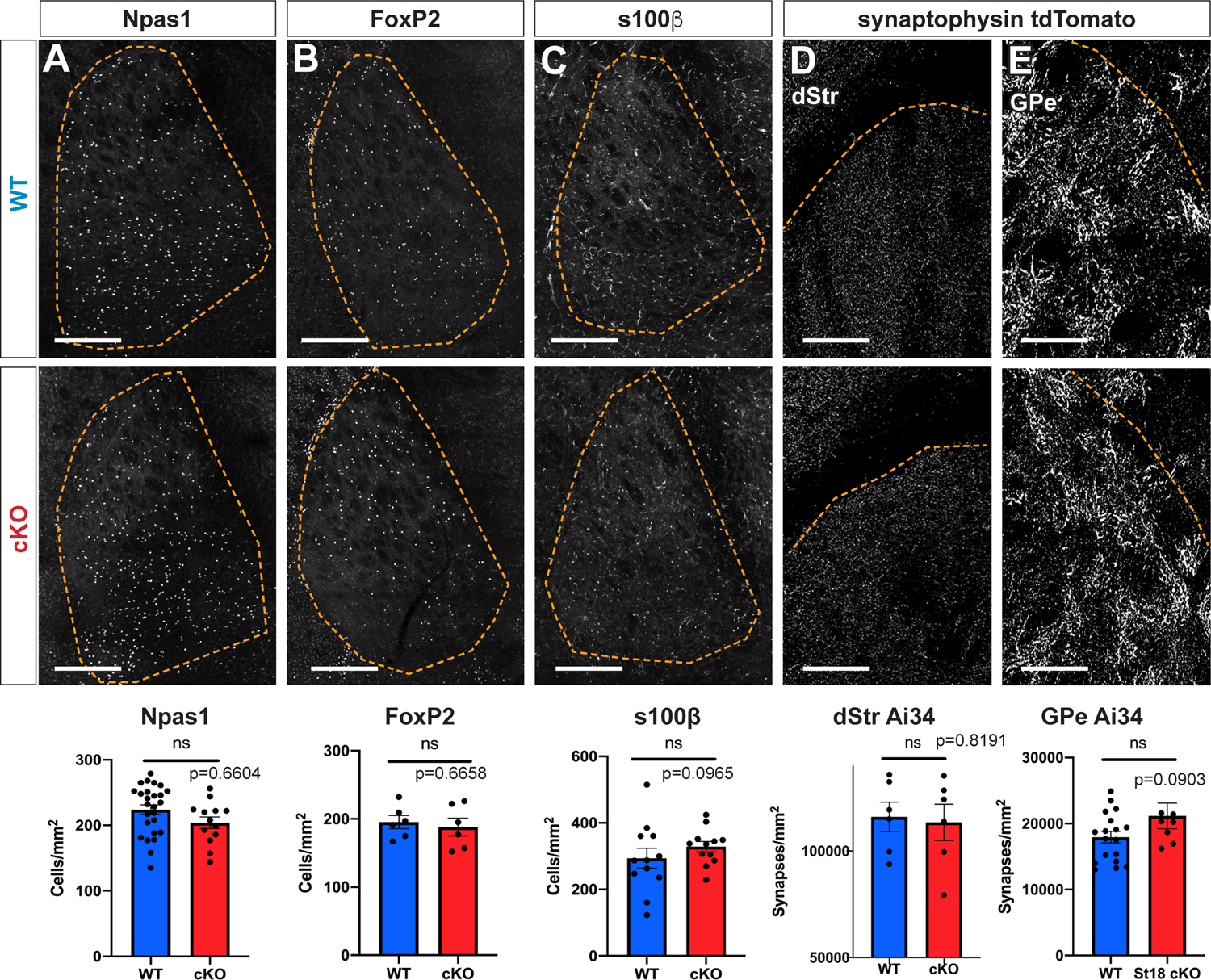
St18 cKO does not cause any change in the arkypallidal neuron population in the GPe. (A) Quantification of Npas1+ neurons (Unpaired t-test; N= 26 WT and 13 St18 cKO). (B) Quantification of FoxP2+ neurons (Unpaired t-test; N= 6 WT and 6 St18 cKO). (C) Quantification of s100b glia cells (Unpaired t-test; N= 12 WT and 12 St18 cKO). (D) Quantification of Ai34 synaptic puncta density in the dStr (Unpaired t-test; N= 6 WT and 6 St18 cKO). (E) Quantification of Ai34 synaptic puncta density in the GPe (Unpaired t-test; N= 18 WT and 8 St18 cKO).

**Figure S7.**
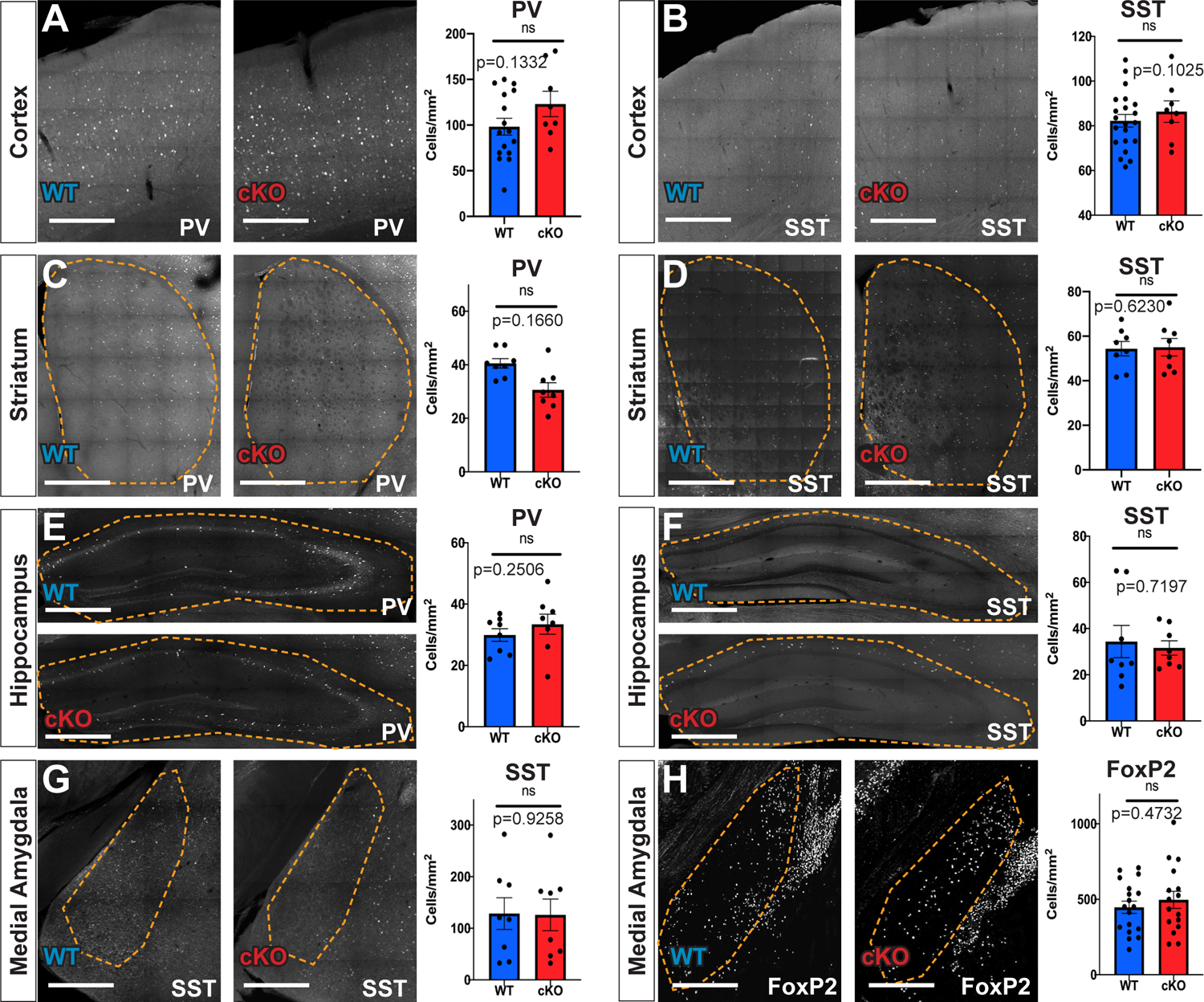
Conditional St18 KO does not affect MGE lineages of the cortex, striatum, hippocampus, and MeA. Quantification of (A) PV+ (N= 16 WT and 8 St18 cKO) and (B) SST+ (N= 21 WT and 8 St18 cKO) neurons in the cortex. Quantification of (C) PV+ (N= 8 WT and 8 St18 cKO) and (D) SST+ (N= 8 WT and 8 St18 cKO) neurons in the Striatum. Quantification of (E) PV+ (N= 8 WT and 8 St18 cKO) and (F) SST+ (N= 8 WT and 8 St18 cKO) neurons in the hippocampus. Quantification of (G) SST+ (N= 8 WT and 8 St18 cKO) and (H) FoxP2+ (N= 18 WT and 16 St18 cKO) neurons in the MeA. (Unpaired t-test).

**Figure S8.**
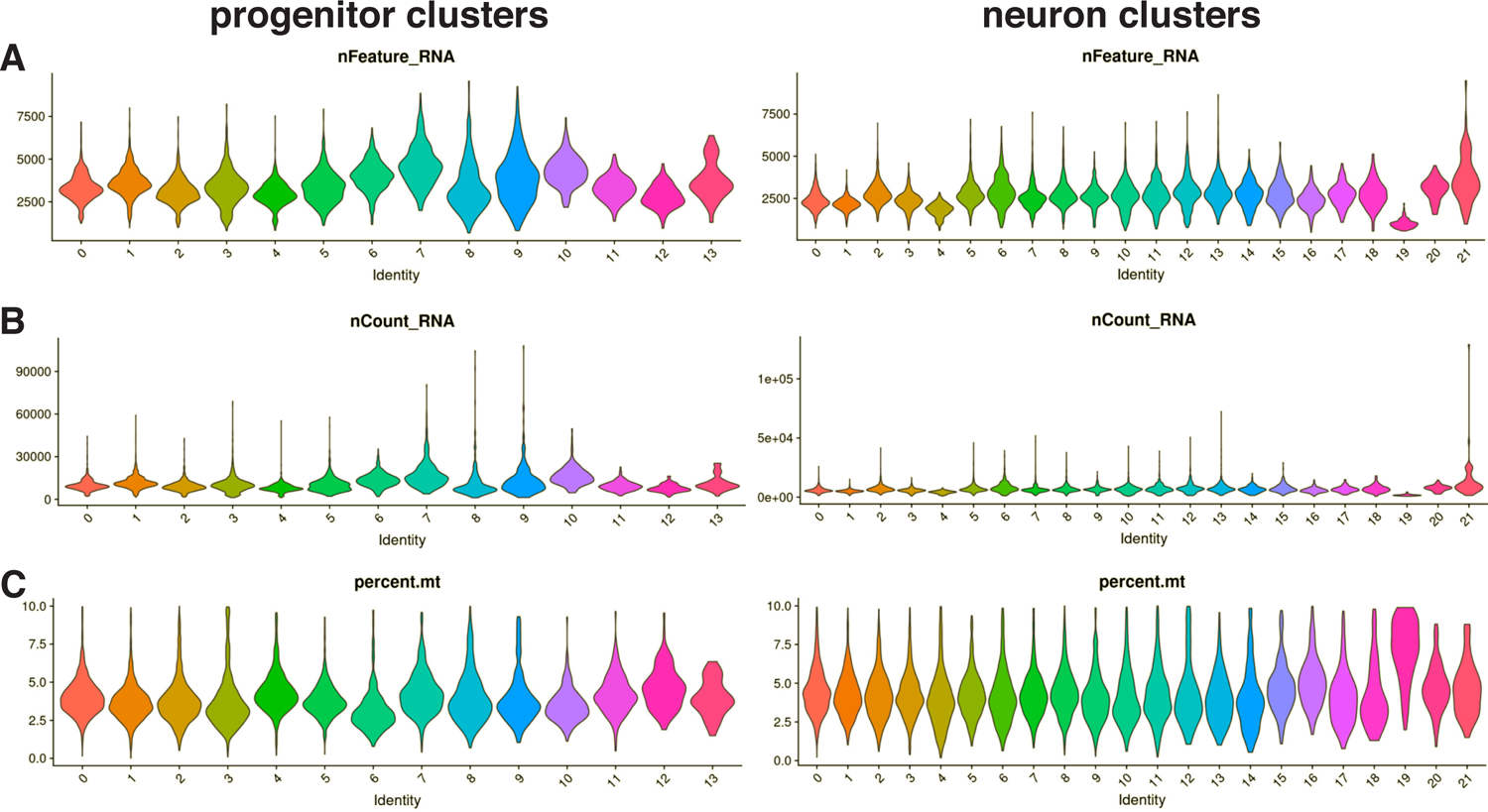
Quality control of single cell RNA-seq dataset. Violin plots showing the distribution of (A) unique genes detected per cell, (B) Unique molecular identifier (UMI) counts per cell, and (C) percent of UMIs mapped to the mitochondrial genome, grouped by cluster. Left, progenitor clusters; right, neuron clusters. Cells with low unique genes detected or high percentage of mitochondrial reads were removed prior to clustering.

**Figure S9.**
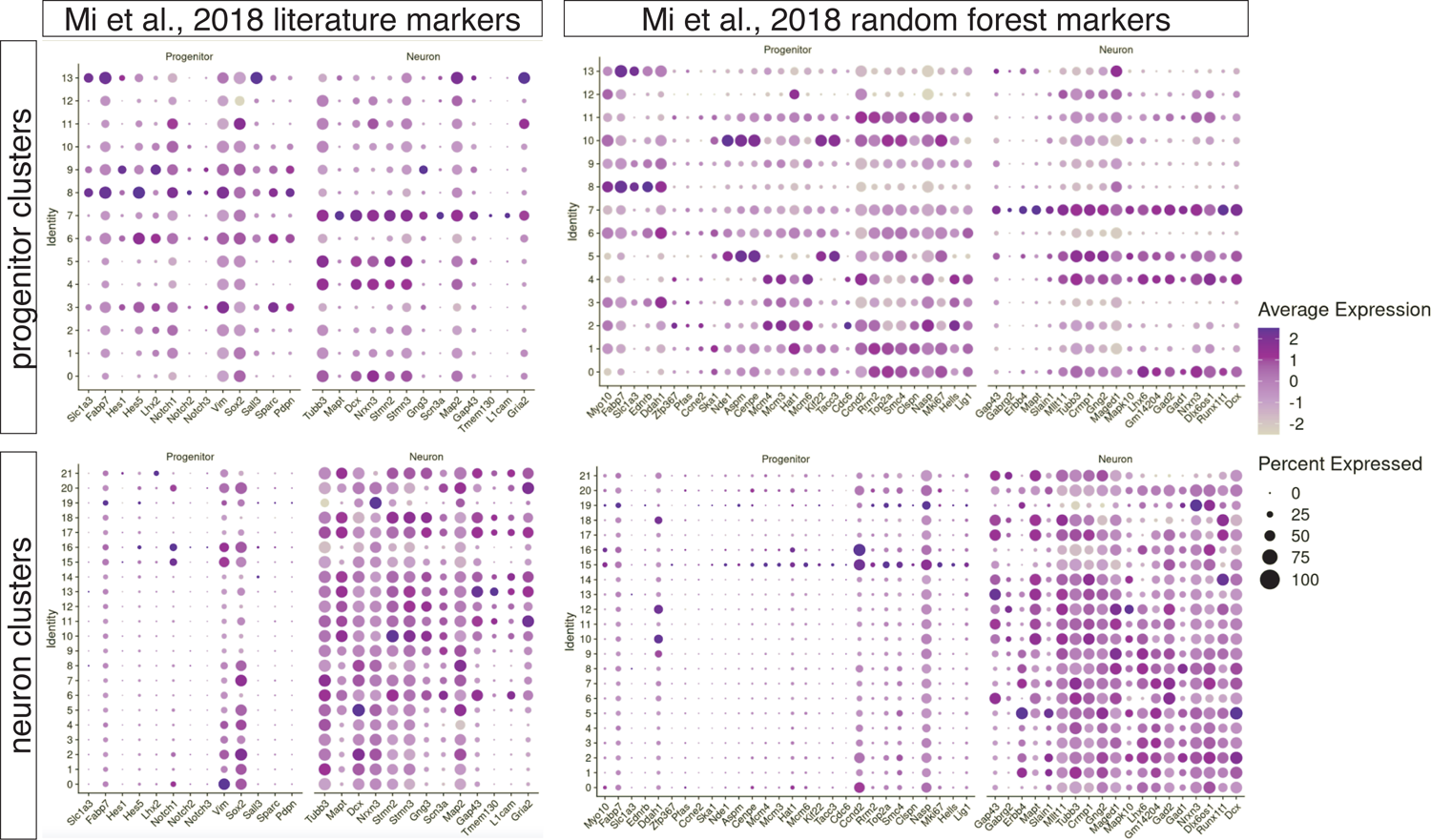
Classification of neuronal and progenitor clusters. Dot plots showing scaled average expression of genes from (Mi et al. 2018) distinguishing neuron and progenitor clusters. Left column, established neuron and progenitor markers from literature. Right column, neuron and progenitor markers as selected by a random forest classifier. Top row, expression of clusters classified as progenitor. Bottom row, expression of clusters classified as neuronal. Clusters were assigned to the class whose markers show the greatest average expression.

**Figure S10.**
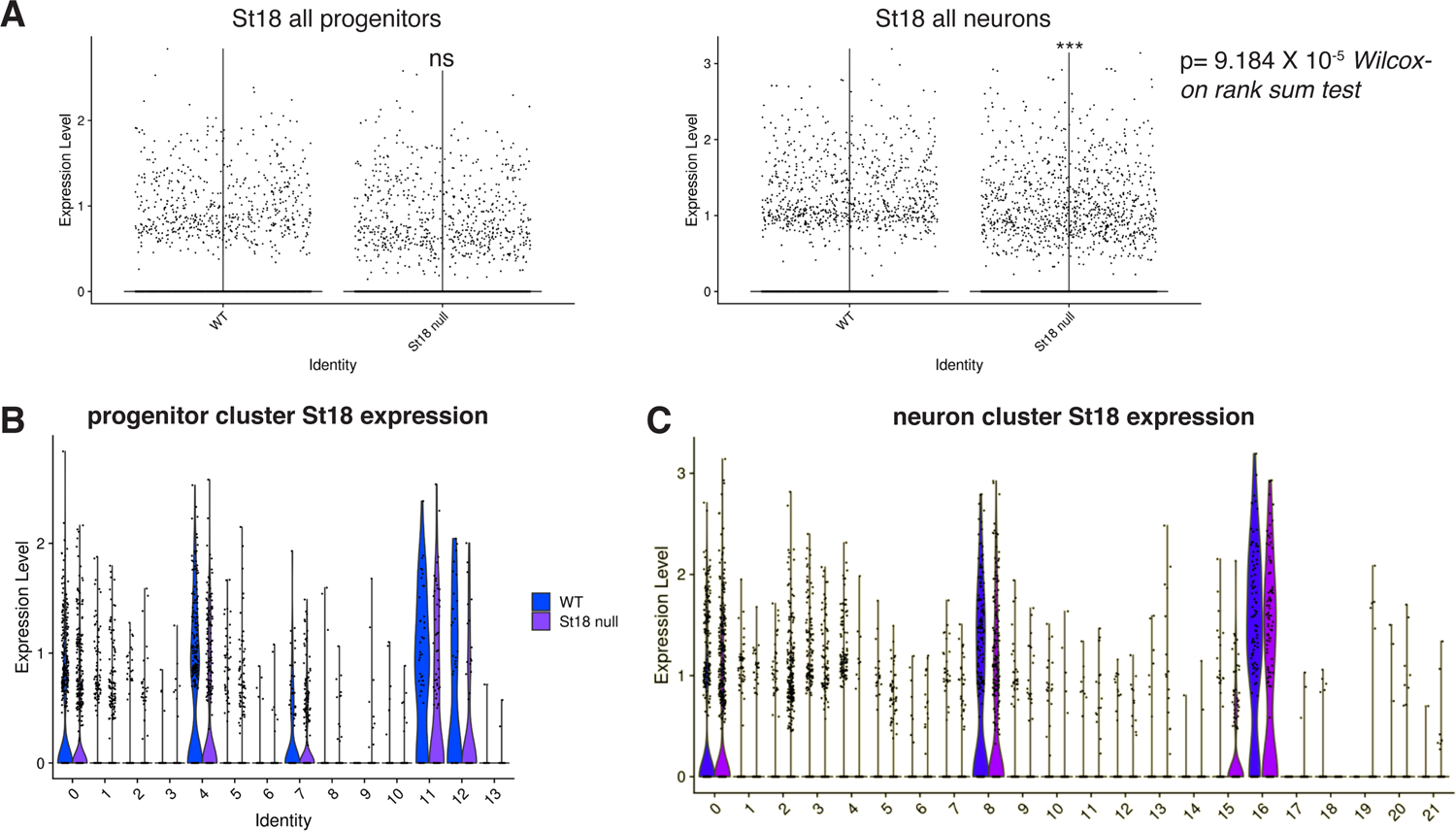
Expression of St18 in single cell RNA-seq. (A) Violin plot showing log-normalized expression of St18 in WT and St18 null progenitors (left) and neurons (right). (B,C) Violin plots of log-normalized expression in WT and St18 null progenitors (B) or neurons (C), grouped by cluster.

**Figure S11.**
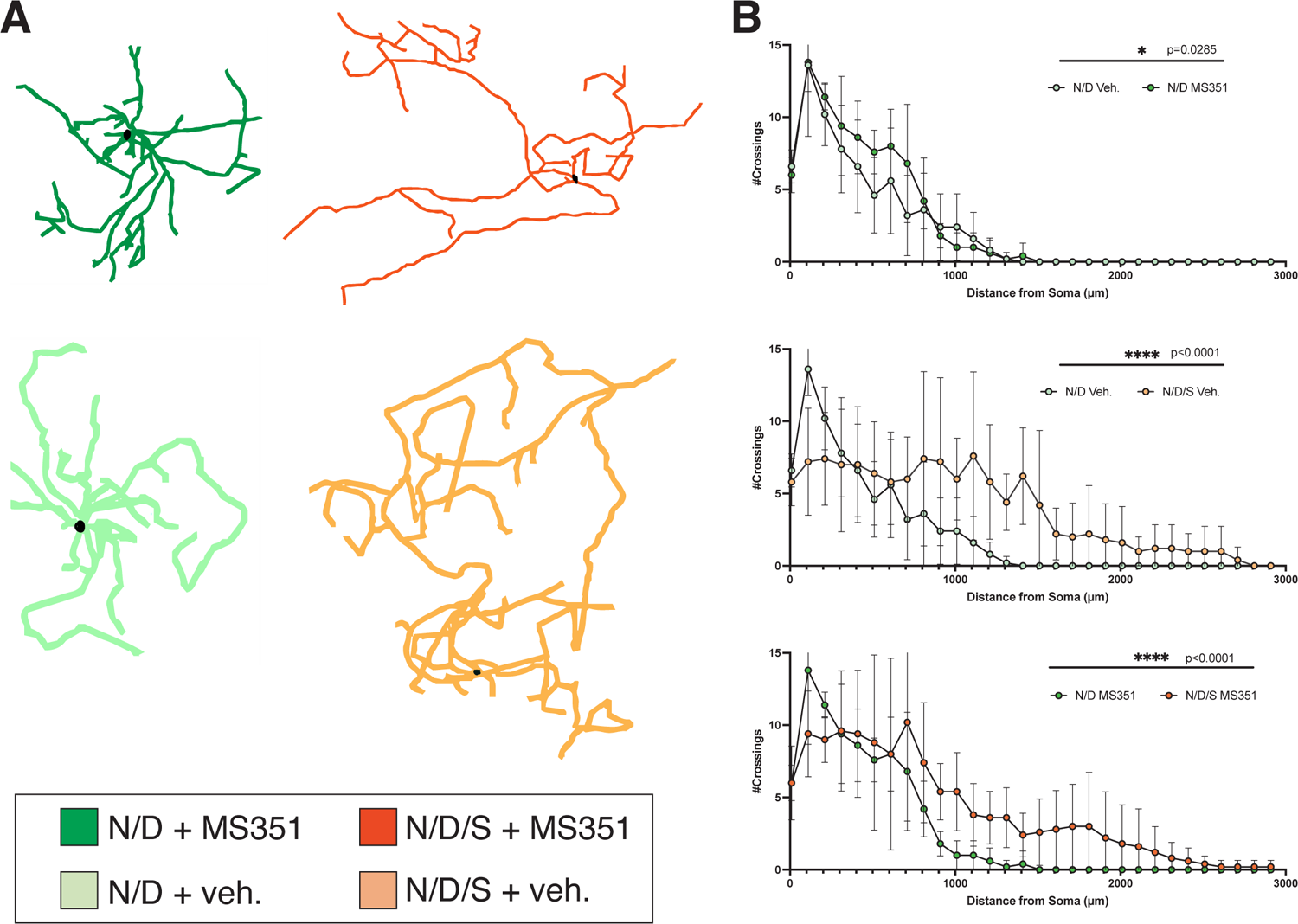
MS351 administration does not affect morphology of N/D and N/D/S neurons. (A) Representative morphological reconstructions of N/D neurons and N/D/S neurons treated with MS351 or DMSO vehicle. (B) Sholl analysis of (top) N/D vehicle vs. N/D MS351; (middle) N/D vehicle vs. N/D/S vehicle; (bottom) N/D MS351 vs. N/D/S MS351. Scholl analyzed by 2-way ANOVA.

## Materials and Methods

### Derivation of N/D and N/D/S lines

We derived a mouse parental ES cell line from a Dlx6a^Cre^ (Jackson Laboratory Stock No. 008199); Ai9 (Jackson Laboratory Stock No. 007909) e3.5 pre-implantation embryo using previously described methods^1^. To this parental line, we stably transfected 2 transgenes, Nestin-Nkx2-1-IRES-tTA^2^S and either TRE-Dlx2 (N/D) or TRE-Dlx2-St18 (N/D/S), as previously described^2^. Briefly, constructs were linearized and introduced into low-passage parental ES cells on a Nucleofector (Lonza) using the mouse ES cell nucleofection kit (Lonza) using protocol “A24”. Cells were plated at 1:20 density on gelatinized 10cm plates and grown in ES cell media for 48 hours. After this time, hygromycin was added to the media for clone selection. Following 1 week of drug selection, ES cell colonies were clonally selected and expanded on gelatinized 96-well plates and expanded to two plates. One plate was cryopreserved at −80^0^C while the other was used for genotyping. Clones were screened by PCR for presence of transgenes. Clones were then expanded and differentiated to screen transgene function before use in experiments. ES cell cultures were maintained and screened using standard practices and protocols^3^ and routinely tested for mycoplasma.

### Specification of ESCs into MGE progenitors

Differentiation of ES cells to become MGE lineage neurons was adapted from protocols previously described^2, 4^. Briefly, ES cell cultures were maintained for two passages on 0.1% gelatin (Millipore) using modified Dulbecco’s Modified Eagle Medium (Gibco), 15% heat-inactivated FBS (HyClone), 1x Modified Eagle Medium Non Essential Amino Acids (Gibco), 1x Sodium Pyruvate (Gibco), and 1x Glutamax (Gibco), 10uM beta-mercaptoethanol (Fisher Scientific), and 10^4^ U/mL ESGRO LIF (Sigma Millipore). N/D and N/D/S line ESCs were dislodged with 0.025% trypsin/1% EDTA (Gibco), dissociated, and suspended at a density of 50,000 cells/well in a non-TC-treated 24 well plate in 800uL of pre-mixed differentiation media (Glasgow’s Modified Eagle Medium [Gibco], 1x PenStrep [Gibco], 1x Modified Eagle Medium Non Essential Amino Acids [Gibco], 1x Sodium Pyruvate [Gibco], and 1x Glutamax [Gibco], 10uM beta-mercaptoethanol [Gibco], and 4% Knockout Serum Replacement [Gibco]) as described previously^2^. Briefly, cells were neuralized and directed toward telencephalic identity with Wnt-inhibitor XAV-939 (Tocris, 0.1 uM) from days 0-4. Sonic hedgehog agonist (SAG [Tocris], 1uM) is added to ventralize telencephalic EBs from days 4-6. From day 6 onwards, on SAG is present in the medium until EB differentiation is complete on day 12. Media changes occurred on days 4, 6, and 9 during the protocol, and free floating embryoid bodies (EBs) containing differentiating neurons form spontaneously during the course of the protocol.

### Embryoid Body fixation and histology

Embryoid bodies (EBs) from two rows of a 24-well plate were gathered in a 15mL conical tube and spun down at 200xg for 2 minutes, rinsed with PBS and re-spun. EBs were then re-suspended in 1mL of 4% PFA and incubated for 30 minutes at room temperature. Post-fixation, EBs were rinsed in PBS and re-suspend in 10% sucrose solution in PBS and incubated for 30 minutes, then re-suspended in 30% sucrose solution in PBS and incubated for an additional 30 minutes. Using a P200 wide bore pipette tip, EBs were removed from the tube and placed in the base of a plastic cryomold (Fisher Scientific). Sucrose solution is removed carefully with both a pipette and the folded edge of a Kimwipe until EBs were as dry as possible at the base of the mold. Tissue-Tek OCT (Sakura) is added to the mold and placed on dry ice for flash freezing and then stored at −30°C. Cryosectioning was performed on a Leica cryostat at a thickness of 15um. Sections were permeabilized using a 0.01% Triton X-100 in PBS and blocked overnight with a 5% normal donkey serum (NDS)/PBS solution. Primary antibodies for V5 (Invitrogen Cat. R960-25, 1:200) and St18 (1:1000) were incubated in 5% NDS/PBS overnight at 4°C and secondary antibodies (Jackson Immunoresearch, 1:1000) diluted in 0.01% Triton X-100/PBS for 2 hours at room temperature. Sections were cover slipped with Fluoromount-G (Southern Biotech) and stored at 4^0^C prior to imaging with Zeiss epifluourescent microscope at 200x.

### Animals

The conditional *St18* allele was generated by introducing flanking LoxP sites to exons 9 and 10 of *St18*^5^. Cre-mediated recombination causes a frame shift in the coding region and a premature stop codon, resulting in the production of a truncated N-terminal fragment. Using an antibody that recognizes the N-terminus of St18, no signal is detected, likely due to a combination of nonsense mediated mRNA decay and peptide instability^6^. The *St18*^fl/fl^ mouse is on a mixed C57Bl/6-ICR background. To generate the *St18* null, we crossed *St18*^fl/fl^ with a germline Cre driver (E2a-Cre Jackson Laboratory Stock No. 003314) Experimental null animals were generated from *St18* heterozygote breeders, and wildtype littermates were used as controls. For *St18* conditional knockout (cKO) experiments, the *St18*^fl/fl^ was crossed to an MGE specific Cre driver (*Nkx2-1*^Cre^ Jackson Laboratory Stock No. 008661) along with the conditional tdTomato reporter (Ai9 Jackson Laboratory Stock No. 007909) or a synaptophysin tagged fate-mapping allele (Ai34 Jackson Laboratory Stock No. 012570) for forward fate-mapping of the *Nkx2-1* lineage. Similarly, *St18*^fl/+^ breeders were used for *St18* cKO experiments to generate *St18* cKO and heterozygote littermate controls. All animal experiments were conducted in compliance with and pre-approval from Columbia University’s Institutional Animal Care and Use Committee.

### 2D culture and morphometric analysis of ESC derived differentiated neurons

Mouse cortical feeders were prepared from P0-P4 wild-type Swiss-Webster mice. Briefly, cortices were micro-dissected into chilled Hibernate A media (Hibernate A[Gibco], 1% Pen/Strep [Gibco]). 1mL of a Papain solution of approximately 20 Units/mL was prepared with Hibernate A dissection media and ∼1000 units of DNAseI and added to dissected tissue. Cortical tissue is agitated at 37^0^C for 15 minutes. Papain is quenched with 3mL Hibernate A, after which cortical tissue is triturated with a P1000 pipette tip until fully dissociated (8-10 times). Cells were counted, spun at 200xg for 5 minutes, and re-suspended in modified Neurobasal A to a concentration of 6 X10^7^ viable cells per mL, and 50uL cell suspensions were plated per well. In parallel, EBs were dissociated in 1mL Papain solution for 15 minutes, 37°C with agitation. Cells were then diluted approximately 1/1000 in modified Neurobasal A media in order for labeled cells to be plated at low density for single cell morphological analysis. The EB/cortical feeder mix is plated onto m 24-well optical plastic tissue culture plates (Ibidi) pretreated with 20ug/mL poly-D lysine and rinsed thoroughly with sterile water. Neuronal cultures were maintained in a modified Neurobasal A media (500mL Neurobasal A [Gibco], .01M HEPES [Gibco], 1% PenStrep [Gibco], 1X B27 supplement [Gibco] is added on the day of use). Media is changed on the first day following plating, and every other day subsequent to that before cultures were fixed with 4% PFA in PBS after 12 days of culture. Whole wells were imaged at 100x magnification using a Zeiss Apotome tiling microscope. Whole neuronal morphologies were traced using the Neuro-Anatomy Simple Neurite Tracing plugin in ImageJ^7^. Morphometric and Sholl analysis were performed on the traces using Neuro-Anatomy plugin (ImageJ). Sholl analysis is performed at a standard radius of 10mm^8^.

### In vitro migration assay

Individual differentiation day 12 N/D and N/D/S EBs were placed into the center of a m 24-well optical plastic tissue culture plates (Ibidi), using a wide-bore P200 pipette tip. Media was gently removed before EBs were suspended in Matrigel (Corning) droplets (50-100uL) using pre-chilled P1000 pipette tips. EB/droplets were incubated for 30 minutes at 37^0^C. Following incubation, solidified Matrigel droplets containing EBs were suspended in Neurobasal media and cultured for four days until fixation using 4% PFA in PBS. Whole EBs were imaged at 200x using a Zeiss confocal microscope. Migration distance is analyzed using straight line tool in ImageJ, measuring the distance a cell traveled perpendicular to the edge of the EB.

### RNA preparation and sequencing

Day 12 N/D and N/D/S EBs were collected into a 15mL conical tube and spun down at 200xg for 5 minutes. EBs were rinsed with PBS and then suspended in 500uL 1X AccuMax (Gibco) enzyme solution with 10 Units DNAaseI (Thermo) and incubated for 15 minutes at 37^0^C. The enzyme was quenched with 5mL HEPES buffered saline and triturated to a single cell suspension using a P1000 pipette tip. Cells were spun at 300xg for 3 minutes and resuspend the pellet in 500uL HEPES buffered saline. The suspension was filtered with a 70um cell strainer for flow cytometry. tdTomato+ cells were sorted (BD Influx System) directly into Trizol. RNA was purified using an RNA miniprep kit (Zymo) and measured with a Bioanalyzer (Agilent) to ensure an RIN score of 7 or greater. mRNA was enriched using a standard poly-A pull-down prior to library construction with Illumina TruSeq chemistry. The library is sequenced using Illumina NovaSeq 6000 at the Columbia Genome Center. Samples were multiplexed in each lane with yields of paired-end 100bp reads per sample. RTA (Illumina) was used for base calling and bcl2fastq2 (version 2.20) for fastq format conversion. A pseudoalignment was performed to a kallisto index created from the mouse transcriptome using kallisto (0.44.0). We tested for differentially expressed genes using Sleuth, an R package designed for differential gene analysis from kallisto abundance files.

### Single cell RNA sequencing sample preparation

e14.5 embryos generated from a cross of *St18*^+/-^ x *St18*^+/-^ parents. Embryos were collected and MGE was microdissected into chilled Hibernate A solution. During the dissection, embryo DNA was obtained using the HOTSHOT protocol^9^ to allow for rapid PCR genotyping. *St18* null and WT MGE were pooled and dissociated separately. Samples were submitted to the Sulzberger Genome Center Single Cell Core at the Columbia University Medical Center for library construction using 10X Genomics chemistry. Sequencing is run on a 10X Genomics platform, and analysis was performed on 10X Genomics’ Cellranger software version 5.0.1.

### Single cell RNA sequencing data analysis

After alignment and quantification with Cellranger, cells were filtered based on the following criteria: cells with fewer than 200 unique genes detected, or cells with more than 10% of reads mapping to the mitochondrial genome were excluded from further analysis. After QC, all cells from all experiments were aggregated and clustered using the Seurat package in R, with default parameters, except the following: cell cycle differences among dividing cells were regressed and removed (Mi et al., 2018), and clustering was performed on the 49 principal components with a p-value less than 0.01, as determined by jackstraw resampling^10^, with a resolution of 1.4, based on bootstrapped iterative clustering^11^. After clustering, progenitor and neuronal clusters were identified by plotting the expression of marker genes from^12^. Subsequently, progenitors and neurons were re-clustered separately. Each cluster was then assessed for statistically significant enrichment or depletion in the WT versus *St18* (-/-) groups, using a Fisher’s exact test with Bonferroni correction for multiple comparisons. This identified three neuronal clusters with statistically significant differences between the groups in terms of proportions, with the additional constraints of comprising >5% of the total neuronal population and at least two-fold change in proportions between the groups. Differential gene expression among the clusters was performed using a Wilcoxon rank-sum test in the Seurat package.

### *In vivo* tissue preparation and histology

For adult histology, P30-P40 animals older were anesthetized using a cocktail of Ketamine and Xylazine and were sacrificed using a trans-cardiac perfusion of PBS followed by 4% PFA/PBS. Following dissection, brains were post fixed for 24 hours in 4% PFA/PBS at 4^0^C and then mounted in 4% low melt agarose to prep for vibratome sectioning. Brains were sectioned in the coronal plane at a thickness of 50um. Sections were collected in series in 96-well U-bottom plates in an anti-freeze solution (1:1:1 PBS, poly-ethylene glycol, and glycerol) for long term storage at −30°C. For immunohistochemistry, free-floating sections were permeabilized with 0.01%Triton X-100/PBS. Primary antibodies were incubated for three days at 4^0^C in 5% NDS/PBS (PV [ImmunoStar Cat. 24428] 1:1000, Npas1 [Gift from the Chan Lab, Northwestern University] 1:1000, FoxP2 [Gift from the Jessel Lab, Columbia University] 1:1000, Er81 [Gift from the Jessel Lab, Columbia University] 1:32000, SST [Millipore Sigma Cat. MAB354] 1:250, s100β [Millipore Sigma Cat. SAB5600115] 1:1000, Nkx2-1 [Abcam Cat. Ab227652] 1:500), and secondary antibodies (Jackson Immunoresearch). Rat-anti St18 antibody was produced in the as described previous^6^. Briefly, the N-terminus of St18 (aa 60-298) was fused to a maltose binding protein to generate a purified immunogen. Strategic Bio-Solutions (Newark, DE) used the immunogen to generate rat polyclonal anti-St18 antibody.

Sections were mounted by paint brush onto SuperFrost Plus slides in Fluoromount G solution for long term storage. For embryonic histology experiments, e13.5 timed plugs were prepared using *St18* null heterozygous breeders, and pregnant dams were sacrificed using carbon dioxide euthanasia and cervical dislocation prior to embryo dissection. Embryos were immersion fixed in 4% PFA/PBS for 15 minutes, rinsed with PBS, before overnight cryoprotection in 30% sucrose/PBS solution. Following cryoprotection, embryos were mounted in OCT compound and flash frozen on dry ice and stored at −30^0^C prior to sectioning. Cryosectioning was performed on a Leica cryostat at 15 um thickness, and sections were dried at room temperature for 1 hour before freezing at −30^0^C. Immunolabeling was performed following a 20 minute wash in PBS to remove excess OCT. Primary antibodies were incubated overnight at 4^0^C in 5% NDS/PBS (Nkx2-1 [abcam Cat. Ab227652] 1:500, St18, 1:1000, Ki67 [Invitrogen Cat. 14-5698-82] 1:200, Caspase 3 [Novus Cat. 31A1067] 1:500), and secondary antibodies (Jackson Immunoresearch). Sections were cover slipped in Fluoromount G solution for long term storage.

### Imaging and image analysis

All imaging was performed at 200x using either a Zeiss scanning confocal microscope or a Zeiss epifluourescent tiling microscope at specific ROIs based on the Allen Brain Institute Reference Atlas. All adult histology (PV, Npas1, Nkx2-1, Er81, FoxP2, SST, and s100β) and tdTomato analysis was performed following imaging with the Zeiss epifluourescent scope at 200x, and representative images for figures were taken with the Zeiss confocal microscope at 200x. Image analysis was performed blind using the Blind Analysis Tool in ImageJ. Images were post-processed using combination of custom ImageJ macros for maximum z-projection, background subtraction (rolling ball radius=20 microns), brightness and contrast adjustments, thresholding, and particle analysis for semi-automated cell and object counting. All embryonic histology (Nkx2-1, Ki67, Casp3, and St18) was performed following imaging with the Zeiss epifluourescent scope at 200x, and representative images for figures were taken with the Zeiss confocal microscope at 200x. Image analysis was performed blind as with adult tissue. Images were post-processed using a combination of custom ImageJ macros for maximum z-projection, background subtraction (rolling ball radius=5 microns), brightness and contrast adjustments, thresholding, and particle analysis for semi-automated cell and object counting. All Ai34 synaptic puncta image analysis were performed following imaging at 200x on a Zeiss confocal microscope. Images were post-processed using a combination of custom ImageJ macros for Z-stack slice selection, background subtraction (rolling ball radius=2 microns), brightness and contrast adjustments, thresholding. Synaptic puncta were quantified by particle analysis for semi-automated object counting (ImageJ). All imaging cell counts were normalized to the area of the section being counted.

### MS351 Treatment

MS351^13^ [BOC Sciences CAS 472984-79-5]) was reconstituted to a 3M stock concentration in a solution of 50% water and 50% DMSO (Sigma) per manufacturer specifications. Throughout the entire EB differentiation as well as during the migration assay, or when grown in monolayer cultures, the stock MS351 solution was diluted to 1mM or we added an equivalent volume of 50% water/50% DMSO as a control. To verify MS351 efficacy on N/D/S cells, MS351 or vehicle-treated day 12 N/D/S EBs were gathered and spun at 300xg to form pellet, washed with 1x PBS and re-spun at 300xg We then suspended pellet in Trizol (Invitrogen Cat. 15596026) and lysed the cells with vigorous trituration and incubated the cells for 5 minutes at room temperature. We then added 200uL of chloroform (Sigma) and vortexed the lysed cells for 15 seconds before incubating the cells at room temperature for another 3 minutes. Next, the cells were spun at 10,000xg, and we removed the clear, aqueous phase from the Trizol reagent and proceeded with RNA mini-prep using the RNeasy Micro Kit (Qiagen Cat. 74004). We next took eluted RNA and proceeded with reverse transcription to create cDNA libraries for RT PCR. Briefly, we add 200ng of eluted RNA to a mix of water, 1mM dNTP Solution Mix (New England Biosciences Cat. N0447L), and 5uM Oligo(dT)20 Primer (Invitrogen Cat. 18418020) and incubated at 65^0^C for 5 minutes. Next, the mixture is cooled on ice for 1 minute before adding reagents for reverse transcription from the SuperScript III Reverse Transcriptase Kit (Invitrogen Cat. 18080093) along with 1 unit of RNaseOUT Recombinant RNase Inhibitor (Invitrogen Cat. 10777019) and 1 mM MgCl2 (Thermo Fisher Scientific Cat. R0971). This mixture was then incubated at 50^0^C for 50 minutes before being inactivated at 85^0^C for five minutes, completing cDNA synthesis. We used the following primer sequences for p16^INK4a^ (also known as Cdkn2A) (Fwd: CAACGACCGAATAGTTACG Rvs: CAGCTCCTCAGCCAGGTC) GAPDH (Fwd: GAGTCCACTGGCGTCTTC Rvs: GGGGTGCTAAGCAGTTGGT) for amplification of 100ng of cDNA template from N/D/S cells either treated with MS351 or vehicle using Power SYBR Green Master Mix (Thermo Fisher Scientific Cat. 4368577) qRT PCR protocol^14^. qRT PCR reaction was run on a QuantStudio(TM) 7 Flex System in a 96 well format with associated software performing analysis (Applied Biosystems Cat. 4485701).

### RNAscope

e13.5 timed plugs were dissected and immersion fixed with 4% PFA/PBS for 15 minutes and rinsed with PBS before overnight cryoprotection in 30% sucrose/PBS solution. Following cryoprotection, embryos were mounted in OCT compound and flash frozen on dry ice and stored at −30^0^C prior to sectioning. Cryosectioning was performed on a Leica cryostat at 15 um thickness in the coronal plane, and sections were dried at room temperature for 1 hour before storage at −30^0^C. The RNAscope assay for fixed-frozen tissue was performed on the sections using the ACD RNAscope Fluorescent Multiplex Reagent Kit (ACD, Cat. 320293) and a catalog probe for St18 transcript (Cat. 443271). Sections were imaged on a Zeiss confocal microscope at 200x.

